# Retromer Opposes Opioid-Induced Downregulation of the Mu Opioid Receptor

**DOI:** 10.1101/2024.12.02.626482

**Authors:** Aleksandra Dagunts, Hayden Adoff, Brandon Novy, Monica De Maria, Braden T Lobingier

## Abstract

The mu opioid receptor (MOR) is protected from opioid-induced trafficking to lysosomes and proteolytic downregulation by its ability to access the endosomal recycling pathway through its C-terminal recycling motif, LENL. MOR sorting towards the lysosome results in downregulation of opioid signaling while recycling of MOR to the plasma membrane preserves signaling function. However, the mechanisms by which LENL promotes MOR recycling are unknown, and this sequence does not match any known consensus recycling motif. Here we took a functional genomics approach with a comparative genome-wide screen design to identify genes which control opioid receptor expression and downregulation. We identified 146 hits including all three subunits of the endosomal Retromer complex. We show that the LENL motif in MOR is a novel Retromer recycling motif and that LENL is a necessary, sufficient, and conserved mechanism to give MOR access to the Retromer recycling pathway and protect MOR from agonist-induced downregulation to multiple clinically relevant opioids including fentanyl and methadone.

## INTRODUCTION

G protein-coupled receptor (GPCR) signaling is responsible for many physiological responses to hormones and neurotransmitters, and regulation of GPCR signaling is critical for cellular homeostasis (Leysen et al., 2021; Zhang et al., 2024; Tse and Wong, 2019). One mechanism for GPCR regulation involves agonist-induced receptor trafficking (Lobingier and von Zastrow, 2019). Agonist-bound GPCRs are endocytosed and trafficked to endosomes where they are sorted to lysosomes for proteolytic downregulation. Consequently, GPCR downregulation following prolonged or repeated agonist stimulation can cause a loss of cellular responsiveness to agonist when lysosomal proteolysis outpaces new receptor synthesis (Stafford et al., 2001; Alvarez et al., 2002; Doss et al., 1981; Heck and Bylund, 1998). However, some types of GPCRs can resist downregulation following agonist addition. These receptors contain recycling motifs in their cytoplasmic facing C-terminal tails that can be recognized by endosomal recycling complexes (von Zastrow, 2001; Irannejad and Lobingier, 2022). These complexes sort GPCRs into endosomal tubules and return them to the plasma membrane, thereby preventing GPCR degradation in the lysosome. Consequently, recycling of GPCRs from endosomes opposes GPCR trafficking to the lysosome and thus acts a brake to slow the process of agonist-induced downregulation and preserve cellular responsiveness to GPCR agonists (Law et al., 2000; von Zastrow, 2001; Bowman and Puthenveedu, 2015).

The best-characterized GPCR endosomal recycling pathway functions through the protein sorting nexin 27 (SNX27), which recognizes and binds class I PDZ binding motifs in the GPCR carboxy terminal tail ([S/T]xΦ-COOH, where Φ is hydrophobic and x can be any amino acid) and links the receptor to additional recycling complexes (Lauffer et al., 2010; He et al., 2006; Gavarini et al., 2006; Romero et al., 2011; Temkin et al., 2011) (**Supp. Fig. 1A**). However, only about 4% of GPCRs contain a class I PDZ motif (Irannejad and Lobingier, 2022; Marchese et al., 2008), and several recycling motifs have been identified in recycling GPCRs that do not match any known consensus recycling motif (Thompson et al., 2014; Olsen et al., 2019; Vargas and Von Zastrow, 2004; Kishi et al., 2001), suggesting the existence of additional mechanisms for GPCR recycling.

One example of a recycling GPCR with a non-consensus recycling motif is the mu opioid receptor (MOR), which mediates the physiological effects of opioids (Sora et al., 1997; Matthes et al., 1996; Kieffer and Gavériaux-Ruff, 2002). Prior work identified a novel type of recycling motif in the final seventeen amino acids of the MOR C-terminal tail - defined by the sequence LENL - that was necessary and sufficient to protect opioid receptors (OR) from agonist-induced downregulation by promoting OR recycling from endosomes (Tanowitz and von Zastrow, 2003). Extensive mutational analysis defined an LxxL core to the motif and showed that the two leucines in the LENL motif, but not surrounding residues, were critical for recycling. However, the LENL sequence does not resemble any known consensus motif recognized by endosomal recycling complexes (**Supp. Fig. 1A**) (Yong et al., 2022). Thus, LENL is an example of a non-consensus recycling motif and the mechanism for MOR recycling is unknown (Bowman and Puthenveedu, 2015; Chen et al., 2023). Uncovering the mechanism that mediates MOR recycling and downregulation has potential clinical relevance, as endocytosis and post-endocytic trafficking of MOR have been implicated in the development of pharmacological tolerance to opioids (Stafford et al., 2001; Kliewer et al., 2019; Kim et al., 2008; Enquist et al., 2012, 2011), a phenomenon which limits the utility of opioids in chronic pain treatment (Buntin-Mushock et al., 2005; Morgan and Christie, 2011) and contributes to opioid toxicity (Strang et al., 2003; Waddell et al., 2020).

We recently developed a chemical biology and functional genomics platform for unbiased identification of proteins involved in agonist-induced GPCR downregulation at the lysosome (Novy et al., 2024). This approach uses a highly sensitive fluorogenic biosensor for GPCR expression in cells, called GPCR-APEX2/Amplex Ultra Red (AUR), to measure loss of GPCR expression due to agonist-induced trafficking of the GPCR-APEX2 genetic fusion to the lysosome. This method leverages the fact that engineered ascorbate peroxidase 2 (APEX2) enzymatic activity is quenched in the lumen of lysosomes by protease activity and low pH, providing a rapid and highly sensitive readout of agonist-induced GPCR downregulation (Novy et al., 2024). Additionally, this method is compatible with pooled genetic screens, allowing for genome-wide interrogation using approaches like CRISPR interference (CRISPRi) to identify genes involved in GPCR expression and trafficking (Novy et al., 2024).

Here, we demonstrate that the GPCR-APEX2/AUR downregulation assay can be used to identify factors that oppose GPCR downregulation by promoting GPCR recycling. This discovery allowed us to conduct a genome-wide CRISPRi screen to identify the proteins controlling GPCR recycling via the MOR recycling motif LENL. This screen revealed all three components of the heterotrimeric recycling complex Retromer as essential to recycling via the LENL motif. We show that this novel mechanism for accessing the Retromer recycling pathway is necessary, sufficient, and conserved and functions to oppose lysosomal downregulation of MOR in response to opioids including fentanyl and methadone.

## RESULTS

### The GPCR-APEX2/AUR downregulation assay captures changes in receptor recycling

We have previously developed a highly sensitive biosensor for GPCR trafficking to the lysosome called GPCR-APEX/AUR, and used this approach to identify novel endosomal genes promoting opioid-induced downregulation of DOR, a poorly recycling GPCR lacking recycling motifs (Novy et al., 2024). We reasoned, however, that this assay would also be capable of capturing the inverse process and identifying genes which oppose GPCR trafficking to the lysosome by promoting receptor recycling from endosomes. If correct, the GPCR-APEX2/AUR assay could be combined with a genome-wide screen to identify genes which mediate MOR recycling through its non-consensus LENL motif. (**Supp. Fig. 1A**)

To test the hypothesis that the GPCR-APEX2/AUR assay could capture changes in downregulation due to presence or absence of a recycling motif, we created HEK293 cells that stably expressed MOR(WT) (flagMOR_WT_-linker-APEX2) under the low expressing UBC promoter. We also made a non-recycling MOR variant, MOR(2Ala), in which the two leucines of the LENL motif were mutated to alanines (flagMOR_2Ala_-linker-APEX2) (**Fig. 1A**). We then examined agonist-induced trafficking of these constructs on two different timescales to capture MOR recycling (short-term, 1 hour after agonist addition) and MOR lysosomal downregulation (long-term, over 6 hours after agonist addition) (**Fig 1A**).

**Figure 1.**
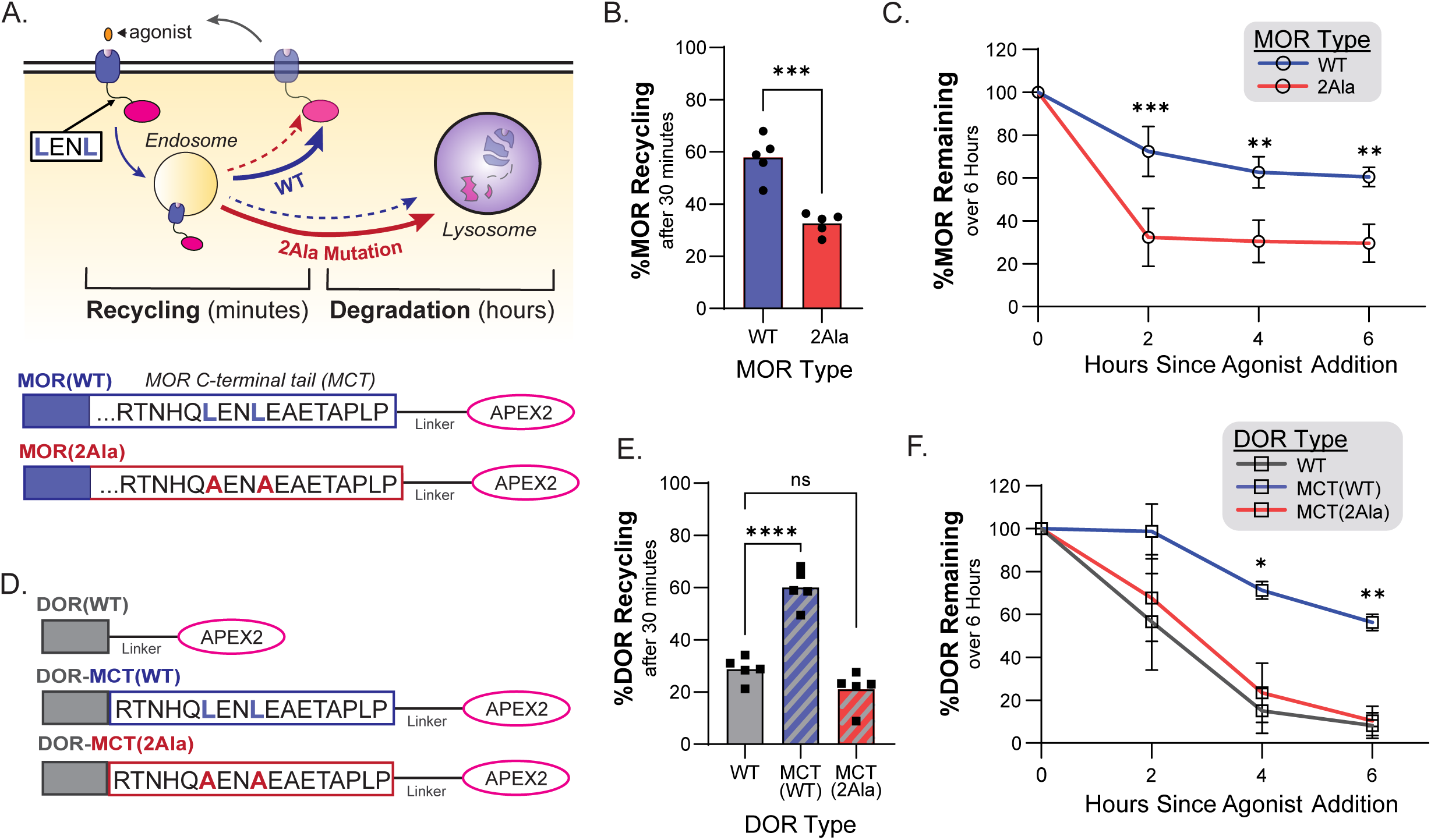
The GPCR-APEX2/AUR downregulation assay captures changes in receptor recycling. **A.** Construct design of APEX-tagged MOR(WT) and MOR(2Ala) and schematic depicting their expected cellular trafficking over time, including major (solid) and minor (dotted) pathways. **B.** Percent recycling of internalized receptors following 30 minutes of treatment with 10μM DAMGO followed by 30 minutes of treatment with 10μM naloxone measured by surface receptor staining (n=5, two-tailed paired t-test, p=0.0004). **C.** Percent receptor remaining following treatment with 10μM DAMGO for 0, 2, 4, or 6 hours normalized to no agonist treatment, measured with the GPCR-APEX2/AUR assay (n=3, 1way repeated measures ANOVA with Sidak’s multiple comparisons correction, p=0.0199 for receptor type effects, p<0.0001 for time effects, p=0.0008, 0.0025, and 0.0031 for WT vs 2A at the 2, 4, and 6 hour timepoints respectively). Error bars denotes S.D. **D.** Construct design of APEX-tagged DOR(WT), DOR-MCT(WT), and DOR-MCT(2Ala). **E.** Percent recycling of internalized receptors following 30 minutes of treatment with 10μM DADLE followed by 30 minutes of treatment with 10μM naloxone measured by surface receptor staining (n=5, repeated measures ANOVA with Dunnett’s multiple comparisons correction, p<0.0001 for WT vs. MCT(WT), p=0.1863 for WT vs. MCT(2Ala)). **F.** Percent receptor remaining following treatment with 10μM DADLE for 0, 2, 4, or 6 hours normalized to no agonist treatment, measured with the GPCR-APEX2/AUR assay (n=3, 2way repeated measures ANOVA with Dunnett’s multiple comparisons correction, p=0.0120 for receptor type effects, p=0.0043 for time effects, p=0.2470, 0.0160, and 0.0011 for WT vs. MCT(WT) at 2, 4, and 6 hours respectively, and p=0.5847, 0.3695, and 0.6600 for WT vs. MCT(2Ala) at 2, 4, and 6 hours respectively. Error bars denote S.D.

To measure opioid-induced MOR internalization and recycling, we used DAMGO, an opioid peptide agonist derived from MOR’s endogenous agonist enkephalin, in field-standard flow cytometry assays (Chen et al., 2023; Knisely et al., 2008; Lau et al., 2011) to quantify surface MOR expression with antibody staining. Consistent with previous observations (Tanowitz and von Zastrow, 2003), MOR(WT) could recycle from endosomes, but MOR(2Ala) recycling was strongly reduced (57.82% vs 32.58%, p=0.0004) (**Fig. 1B**, **Supp. Fig. 1B&C**). There was also a slight increase in the proportion of MOR(2Ala) that was internalized following agonist stimulation, which is consistent with reduced recycling during the 30 minutes of agonist stimulation resulting in a larger portion of the receptor population remaining in cells (**Supp. Fig. 1D**). These results demonstrate that LENL-mediated recycling of MOR is maintained in the presence of the APEX2 tag and are consistent with previous observations showing APEX2-tagging MOR does not disrupt its trafficking or signaling (Lobingier et al., 2017; Polacco et al., 2024; Novy et al., 2024).

We next asked if the GPCR-APEX2/AUR assay could detect an increased rate in opioid-induced MOR lysosomal downregulation due to a mutated recycling motif. Over six hours of agonist treatment, MOR(WT) showed a small amount of downregulation that was greatly increased in the MOR(2Ala) mutant (29.52% receptor loss compared to 60.46% after six hours of agonist treatment, p=0.0199 for receptor type effects) (**Fig. 1C**). This finding demonstrates that the GPCR-APEX2/AUR assay could capture an enhanced rate of agonist-induced downregulation due to disruption of the LENL recycling motif in MOR.

Next, we tested whether the GPCR-APEX2/AUR assay could capture a reduced rate of agonist-induced downregulation in a gain-of-function experiment where the LENL recycling motif was grafted onto a poorly recycling GPCR. We selected DOR as it broadly similar to MOR except that, unlike MOR, DOR lacks a recycling motif and is efficiently targeted to lysosomes upon opioid stimulation (Tanowitz and von Zastrow, 2003; Tanowitz and Von Zastrow, 2002; Milan-Lobo and Whistler, 2011). We created HEK293 cells that stably expressed either DOR(WT) (flagDOR-linker-APEX2) or DOR tagged at its C-terminus with the final 17 amino acids of MOR C-tail (MCT) containing either the functional (flagDOR_MCT_-linker-APEX2) or mutated (flagDOR_MCT(2Ala)_-linker-APEX2) LENL recycling motif (**Fig. 1D**).

First, we examined DOR recycling following stimulation with the opioid peptide DADLE and found that DOR with a functional LENL motif, but not a mutated motif, showed enhanced recycling (28.35%, 60.00%, 22.13% recycling for DOR(WT), DOR-MCT(WT), and DOR-MCT(2Ala) respectively, p<0.0001 for DOR(WT) vs DOR-MCT(WT), p=0.1863 for DOR(WT) vs DOR-MCT(2Ala)) (**Fig. 1E**, **Supp. Fig. 1E&F**). We observed a residual amount of basal recycling for all receptors lacking functional recycling motifs. This finding is consistent with previous observations that GPCRs can also traffic, albeit inefficiently, through of a “bulk flow” recycling pathway (Puthenveedu et al., 2010; Bowman and Puthenveedu, 2015; Bowman et al., 2016; Mayor et al., 1993; Bahouth and Nooh, 2017; Tanowitz and von Zastrow, 2003) which operates in a recycling motif-independent manner to indiscriminately return endosomal cargo to the plasma membrane.

To verify that the functional recycling motif slowed DOR trafficking to the lysosome, we measured DOR downregulation over a six-hour period in the GPCR-APEX2/AUR assay and found 5-fold more DOR remaining in cells expressing DOR(MCT) following chronic agonist stimulation (8.8%, 56.35%, and 10.45% receptor remaining for DOR(WT), DOR-MCT(WT), and DOR-MCT(2Ala) respectively after six hours of agonist treatment, p=0.0120 for receptor type effects) (**Fig. 1F**). Together, these results suggest that the GPCR-APEX2/AUR degradation assay accurately captures the inverse relationship between GPCR recycling and degradation.

### Retromer acts through the LENL recycling motif to oppose agonist-induced opioid receptor downregulation

Since the GPCR-APEX2/AUR assay could capture the effects of GPCR recycling on agonist-induced GPCR downregulation, we considered the possibility that a functional genomic screen could identify the genes which function with the LENL to induce recycling and oppose downregulation. In our previous genome-wide CRISPRi screen focused on the degrading GPCR DOR, we found the GPCR-APEX/AUR assay detected genes involved in the entire GPCR lifecycle including expression, synthesis, and trafficking through both the secretory and endosomal-lysosomal pathways. To specifically focus the current screen on the genes which promote LENL-based recycling, we decided on a comparative screen design in which we would perform a new genome-wide CRISPRi screen with DOR-MCT(WT) and compare the results with our previous CRISPRi screen with DOR(WT). We reasoned that any genes whose knockdown increased GPCR degradation in DOR-MCT(WT) cells, but not in DOR(WT) cells, would be potential candidates for mediating LENL-based recycling.

We utilized the same cell line as before, HEK293-FLP, to generate a reporter line for the CRISPRi screen that stably expressed both SFFV:dCas9-Krab and UBC:DOR_MCT_-APEX2 (**Figure 2A**). We then divided the genome-wide CRISPRi library into three sub-libraries, and the reporter cell line was transduced with each sub-library for an approximate 300-fold sgRNA coverage. Following eight days of gene knockdown, cells were stimulated with agonist to induce receptor degradation and the amount of remaining DOR-MCT(WT) in individual cells was determined with the GPCR-APEX2/AUR assay. Cells were then sorted into the top and bottom quartiles based on their individual fluorescence to identify sgRNAs which decreased GPCR expression (enriched in bottom quartile) or sgRNAs that increased GPCR expression (enriched in top quartile) after agonist exposure (**Fig. 2A**). We hypothesized that genes potentially involved in LENL-based recycling would be enriched in the bottom quartile, since their loss would lead to a decrease in recycling, a subsequent increase in degradation, and loss of overall expression in the cell.

**Figure 2.**
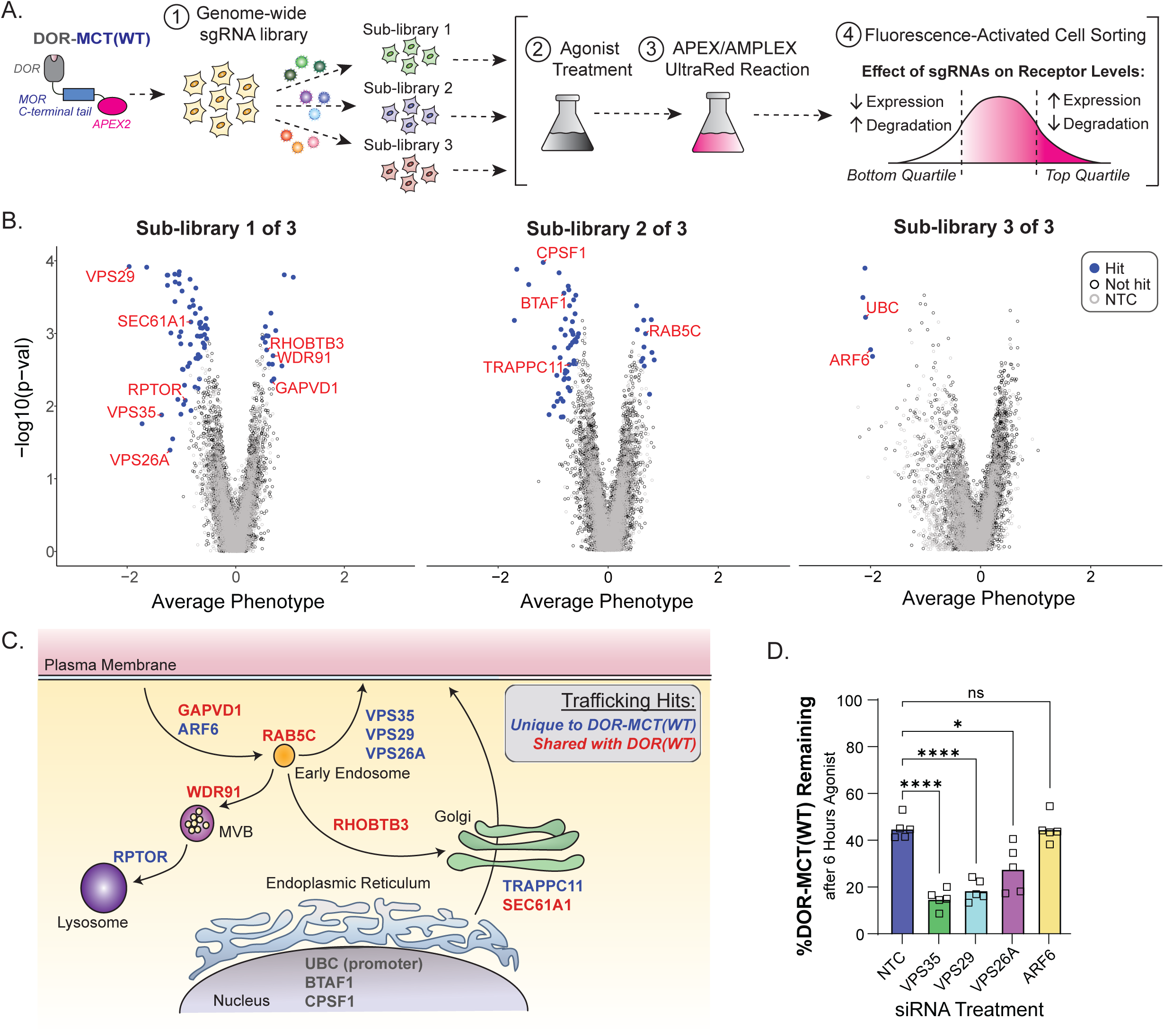
Retromer acts through the LENL recycling motif to oppose lysosomal receptor degradation. **A.** Screen design schematic. **B.** Volcano plots of relative gene enrichment in cells sorted into bottom and top fluorescence quartiles following the AMPLEX reaction divided by sublibrary. Relative sgRNA enrichment between population was analyzed with a Mann-Whitney U-test from n=1 independent experiment. sgRNAs with a false discovery rate of <0.05 are denoted as hits (blue circles), non-targeting control sgRNAs are depicted as open circles, and all other genes are depicted as gray circles. Select hits involved in receptor expression and trafficking are annotated in red. **C.** Cartoon showing proposed location of action for hits involved in receptor expression and trafficking. Hits that were also found in our previous DOR screen are depicted in red, while hits that were unique to DOR-MCT(WT) are depicted in blue. **D.** Percent DOR-MCT(WT) remaining following siRNA knock-down of select hits in cells treated with 6 hours of 10uM DADLE followed by the AMPLEX assay, normalized to no agonist treatment (n=5, 1way repeated measures ANOVA with Dunnett’s multiple comparisons correction, p<0.0001 for NTC vs. VPS35, p<0.0001 for NTC vs. VPS29, p=0.0112 for NTC vs. VPS26A, and p>0.999 for NTC vs. ARF6).

Next generation sequencing revealed 5/5 sgRNAs were found for 88.82% of every gene (and 4/5 were found for 99.02%) suggesting no large bottle necks in the workflow. The screen identified 146 hits (**Fig. 2B**, **Supp. Table 1**). Consistent with our previous screen on DOR(WT), the genome-wide screen on DOR-MCT(WT) identified genes linked to membrane protein expression and trafficking as well as the internal positive control: sgRNAs which target UBC, the promoter driving DOR_MCT_-APEX2 expression (**Fig. 2C**). Demonstrating a conservation of genes for GPCR expression and trafficking, most of these hits (81.5%) were found to be expressed in multiple types of MOR-expressing mouse neurons as well as HEK293 cells (**Supp. Table 2**). Furthermore, 44% of the hits in the DOR-MCT genome-wide screen were also hits in our previous DOR screen, suggesting a broad similarity in genes which regulate expression throughout the GPCR lifecycle (**Supp. Table 1**).

We then sought to identify candidate genes which function with LENL to oppose agonist-induced GPCR downregulation by analyzing hits enriched in the bottom fluorescence quartile that were unique to the DOR-MCT(WT) CRISPRi screen. Several genes matched these criteria, chief among them VPS35, VPS29, and VPS26A, which encode for the three subunits of the endosomal Retromer complex(Seaman, 2021) (**Fig 2B&C, Supp. Table 1**). We were surprised to identify all three Retromer subunits as hits because the consensus recycling motif for Retromer binding, [WYF]x[LMV] (Seaman, 2007; Tabuchi et al., 2010), does not match LENL (**Supp. Fig. 1A**) and is not found in any cytoplasmic-facing residues of MOR. To validate the findings from the CRISPRi screen, we next used siRNAs to individually knock down each Retromer subunit and measured DOR-MCT(WT) downregulation using the GPCR-APEX2/AUR assay. Knockdown of any of the three Retromer subunits significantly increased DOR-MCT(WT) degradation following six hours of agonist treatment. We also examined the small GTPase ARF6, another hit in the screen which was previously shown to be involved in MOR trafficking (Rankovic et al., 2009), but did not observe any effects from ARF6 siRNA knockdown (**Fig. 2D**).

Given the known function of Retromer in promoting membrane protein recycling, we hypothesized that the increased rate of downregulation of DOR-MCT(WT) upon Retromer knockdown was due to a loss in recycling. Measurements of DOR recycling, either in its native sequence (DOR(WT)) or with the LENL motif grafted to its C-terminus (DOR-MCT(WT)), revealed that knockdown of VPS35 reduced DOR-MCT(WT) recycling to DOR-WT levels (52.33% recycling for NTC and 27.33% for VPS35, p=0.0015) (**Supp. Fig. 2A**) but had no effect on DOR(WT) recycling (22.67% for NTC and 20.00% for VPS35, p=0.6122). Together these data suggest that LENL is a novel type of Retromer motif that is sufficient to promote GPCR entry into the endosomal recycling pathway and thereby slow agonist-induced downregulation.

Since LENL represents a potential novel type of Retromer recycling motif, we wanted to determine if other membrane proteins previously linked to Retromer contained the hallmark LxxL pattern of the MOR LENL recycling motif. To accomplish this task, we developed a custom application, Motif Searcher, that can parse through a user-provided list of UniProt Knowledgebase protein identifiers and search for a specified sequence within the whole protein or specific topological stretches of a protein such as cytoplasmic-facing regions. To identify endosomal recycling motifs in the cytoplasmic tails of proteins in the human membrane proteome, we used Motif Searcher to analyze the last 100 amino acids of 2359 verified “cell membrane” proteins. There were 692 unique proteins with a LENL-like LxxL sequence, the majority of which were signaling receptors and transporters (**Supp. Fig. 2B, Supp. Table 3**). As a point of comparison, we used Motif Searcher to perform the same analysis for other previously described recycling motifs which function with Retromer, ESCPE-1, or Retriever: [F/Y/M]x[L/M/V], [D/E][S/T]x<Ι -COOH, <Ιx[F/Y/V]x[F/Y], and NxxY, where Φ is any hydrophobic amino acid (**Supp. Table 3**) (Yong et al., 2022, 2020; Clairfeuille et al., 2016). These motifs were present in 986, 124, 424, and 155 unique proteins respectively and, except for NxxY, were found mostly in signaling receptors and transporters (**Supp. Fig. 2C&D**).

While it is currently unknown what proportion of these computationally identified sequences can act as *bona fide* recycling motifs, several of the LxxL-containing proteins identified in our analysis have been previously linked to Retromer in an unbiased screen for proteins which depend on VPS35 for their surface expression (adenylyl cyclase 9: 1344-LTKL, SLC12A7: 1058-LEVL; PLXNA1: 1867-LAAL, etc) (Steinberg et al., 2013). AC9 is of particular interest because it has been shown to localize to endosomes where it participates in endosomal GPCR signaling (Lazar et al., 2020; Ripoll et al., 2024) We also identified a LENL-like motif in the GLUT4 receptor (LEYL), which has previously been shown to traffic through Retromer-dependent pathways (Yang et al., 2016; Pan et al., 2017). Together, these results demonstrate that Retromer protects opioid receptors with the LENL motif from agonist-induced trafficking to the lysosome, and that LENL, and LENL-like motifs, potentially represent a novel type of Retromer recycling motif.

### The Retromer complex is required for MOR recycling and opposition to opioid-induced lysosomal downregulation

As our results show that Retromer can function with the LENL motif in context of the chimeric DOR-MCT(WT), we next wanted to know if Retromer functioned in the same manner with MOR. We first asked if Retromer was present on MOR-positive endosomes. Using high-resolution Airyscan confocal microscopy, we observed agonist-dependent co-localization of endogenous Retromer subunit VPS35 and MOR(WT) in HEK293 cells stably expressing MOR (**Fig. 3A**, **Supp. Fig. 3A**). Specifically, we often saw Retromer adjacent and partially overlapping with MOR, which is consistent with previous observations of GPCRs which recycling through a SNX27/Retromer-dependent pathway (Puthenveedu et al., 2010; Varandas et al., 2016; Temkin et al., 2011), To quantify this co-localization, we used Imaris image analysis software to render three-dimensional objects based on MOR (FLAG), Retromer (VPS35), or Golgi (GM130) immunofluorescence from the confocal z-stack images (**Supp. Fig. 3B-D**). We then calculated the percentage of every individual MOR object’s volume that overlapped with a Retromer object. We found that on average, each MOR object had a 39.67% volume overlap with a Retromer object, and nearly no overlap with a negative control Golgi surface (1.67%, p=0.0076) (**Fig. 3B**, **Supp. Fig. 3B-E**). Similar results were obtained using a Pearson’s correlation coefficient to quantify MOR/Retromer co-localization (0.18 for VPS35, 0.056 for Golgi, p=0.0160 for VPS35 vs. Golgi) (**Supp. Fig. 3F**). Together, these results demonstrate that Retromer is at the correct place and time to mediate MOR recycling.

**Figure 3.**
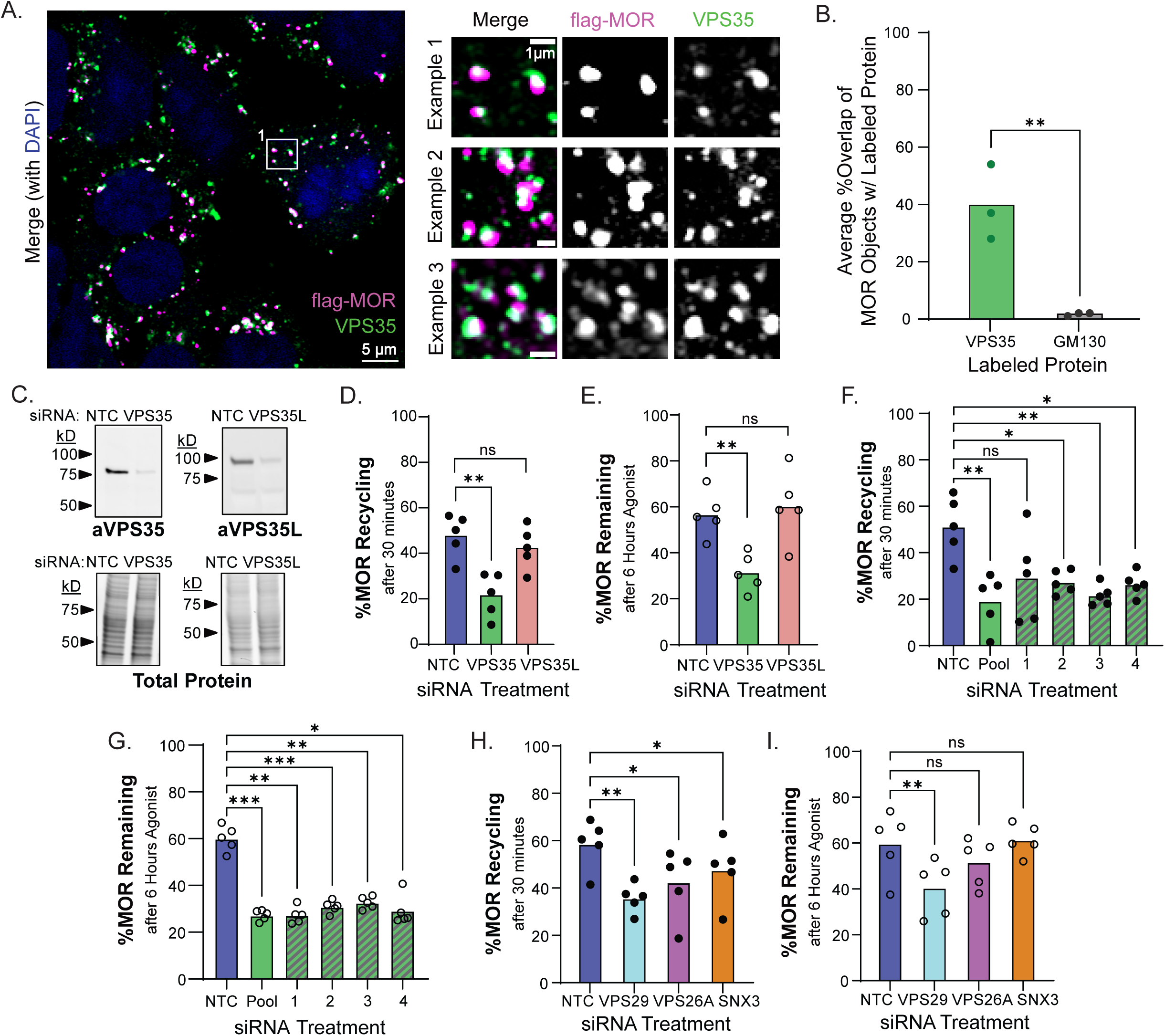
The Retromer complex is required for MOR recycling and resistance to downregulation. **A.** Confocal images of HEK293 cells stably expressing MOR(WT) and treated with 10μM DAMGO, fixed and stained for anti-FLAG (magenta) and anti-VPS35 (green). Representative images shown, n=3. **B.** Average percent overlap of all MOR objects with either VPS35 or GM130 (n=3, unpaired two-tailed t-test, p=0.0076). **C.** Western blot for VPS35 and VPS35L and total protein from HEK293 lysates following treatment with NTC, VPS35, or VPS35L siRNA treatment. Representative blot shown, n=3. **D.** Percent MOR(WT) recycling measured by surface labeling of receptors following treatment with 30 minutes 10μM DAMGO to induce internalization and 30 minutes 10μM naloxone to allow for recycling in cells treated with NTC, VPS35, or VPS35L siRNA (n=5, 1 way repeated measures ANOVA with Dunnett’s multiple comparisons, p=0.0034 for NTC vs VPS35, p=0.3332 for NTC vs. VPS35L). **E.** Percent MOR(WT) remaining following siRNA knock-down of VPS35 or VPS35L and treatment with 6 hours of 10μM DAMGO followed by the AMPLEX/AUR reaction, normalized to no agonist treatment (n=5, 1 way repeated measures ANOVA with Dunnett’s multiple comparisons, p=0.0061 for NTC vs VPS35, p=0.7862 for NTC vs. VPS35L). **F.** Same as D, but following siRNA knock-down of VPS35 with 4 individual siRNAs, or a pool of all four siRNAs (n=5, 1 way repeated measures ANOVA with Dunnett’s multiple comparisons, p=0.0061, 0.1162, 0.0148, 0.0081, 0.0338 for NTC vs. Pool, 1, 2, 3, and 4 respectively). **G.** Same as E, but following siRNA knock-down of VPS35 with 4 individual siRNAs or a pool of all four siRNAs (n=5, 1 way repeated measures ANOVA with Dunnett’s multiple comparisons, p=0.0002, 0.0023, 0.0007, 0.0026, 0.0114 for NTC vs. Pool, 1, 2, 3, and 4 respectively). **H.** Same as D, but after treatment of cells with pooled siRNA against VPS29, VPS26A, or SNX3 (n=5, 1 way repeated measures ANOVA with Dunnett’s multiple comparisons, p=0.0070, 0.0107, and 0.0183 for NTC vs. VPS29, VPS26A, and SNX3 respectively). **I.** Same as E, but after treatment of cells with pooled siRNA against VPS29, VPS26A, or SNX3 (n=5, 1 way repeated measures ANOVA with Dunnett’s multiple comparisons, p=0.0024, 0.0549, and 0.9820 for NTC vs. VPS29, VPS26A, and SNX3 respectively).

Next, we asked if Retromer promotes MOR recycling from endosomes in HEK293 cells stably expressing MOR(WT) following knockdown of the Retromer. As a pathway-specificity control, we also examined recycling following knockdown of a structurally similar but functionally unrelated endosomal recycling complex, Retriever. To knockdown the function of Retromer and Retriever, we targeted the subunits required for assembly of the respective complexes, VPS35 (Retromer) and VPS35L (Retriever). Pooled siRNAs targeting VPS35 or VPS35L resulted in >80% knockdown relative to an NTC (**Fig. 3C**, **Supp. Fig 4A-D**). We found that knockdown of Retromer function, but not Retriever, resulted in loss of MOR recycling (**Fig. 3D**, **Supp. Fig. 4E**). We then examined how loss of Retromer or Retriever function affected opioid induced MOR downregulation. As we observed for MOR recycling, knockdown of VPS35, but not VPS35L, accelerated the rate of MOR downregulation (**Fig. 3E**). To verify the specificity of the siRNA pool targeting VPS35, we examined the individual siRNAs. All siRNAs which made up the pool efficiency caused VPS35 knockdown (**Supp. Fig. 4A&B**), loss of MOR recycling (**Fig. 3F**, **Supp. Fig. 3F**), and enhancement of opioid-induced downregulation (**Fig. 3G**). Together, the data demonstrate Retromer functions to promote MOR recycling from endosomes and thereby slow the rate of opioid-induced MOR downregulation.

We next asked if the other subunits of Retromer are required for MOR recycling and resistance to opioid-induced downregulation. VPS26A is known to bind membrane protein cargos (Lucas et al., 2016) while VPS29 is thought to play a structural and regulatory role (Baños-Mateos et al., 2019; Ye et al., 2020). Like we observed with VPS35, we found that knockdown of VPS29 reduced MOR recycling and increased its downregulation (**Fig 3H&I**, **Supp. Fig. 3G**). We only observed a significant effect from VPS26A knockdown in MOR recycling (although a trend toward affecting MOR downregulation as well), possibly due to genetic compensation from its paralogue, VPS26B (Bugarcic et al., 2011). We also examined SNX3, a known Retromer binding protein important for recruiting Retromer to endosomes (Harrison et al., 2014) and binding cargoes bearing a [WYF]x[LMV] motif (Lucas et al., 2016). We found that SNX3 knockdown reduced MOR recycling from 58.65% to 47.60% (p=0.0183) but had no effect on MOR downregulation (40.18% vs. 38.64%, p=0.9820). Together, these results demonstrate that multiple Retromer subunits, as well as additional Retromer-binding proteins like SNX3, are important in the post-endocytic trafficking of MOR.

### Retromer’s role in MOR recycling is conserved across cell lines

Having shown that Retromer is required for MOR recycling and opposition to opioid-induced MOR downregulation in HEK293 cells, we next asked if its function was conserved in a neuronal derived cell line. We selected the human neuroblastoma SH-SY5Y line because of its neuronal properties and endogenous MOR expression (Kaya et al., 2024; Kazmi and Mishra, 1986). To monitor MOR trafficking, we transduced SH-SY5Y cells to stably express the same flagMOR_WT_-linker-APEX2 construct used in the HEK293 line. Of note, the SH-SY5Y cell line expresses these engineered MORs at an approximately 5-fold lower level than the already low expressing HEK293 line and thus more closely recapitulates endogenous receptor expression (**Supp. Fig. 5A**). As in HEK293 cells, we observed VPS35 both adjacent to and overlapping with MOR-positive endosomes. Both the Pearson’s analysis and the Imaris-based overlap quantification showed that MOR co-localized with Retromer, but not the Golgi, following agonist stimulation in SH-SY5Y cells (**Fig. 4A&B**, **Supp. Fig. 5B-D**). We observed a lower overlap score in SH-SY5Y cells compared to HEK293 cells, which was due to a higher proportion of MOR objects that did not have any overlap with Retromer.

**Figure 4.**
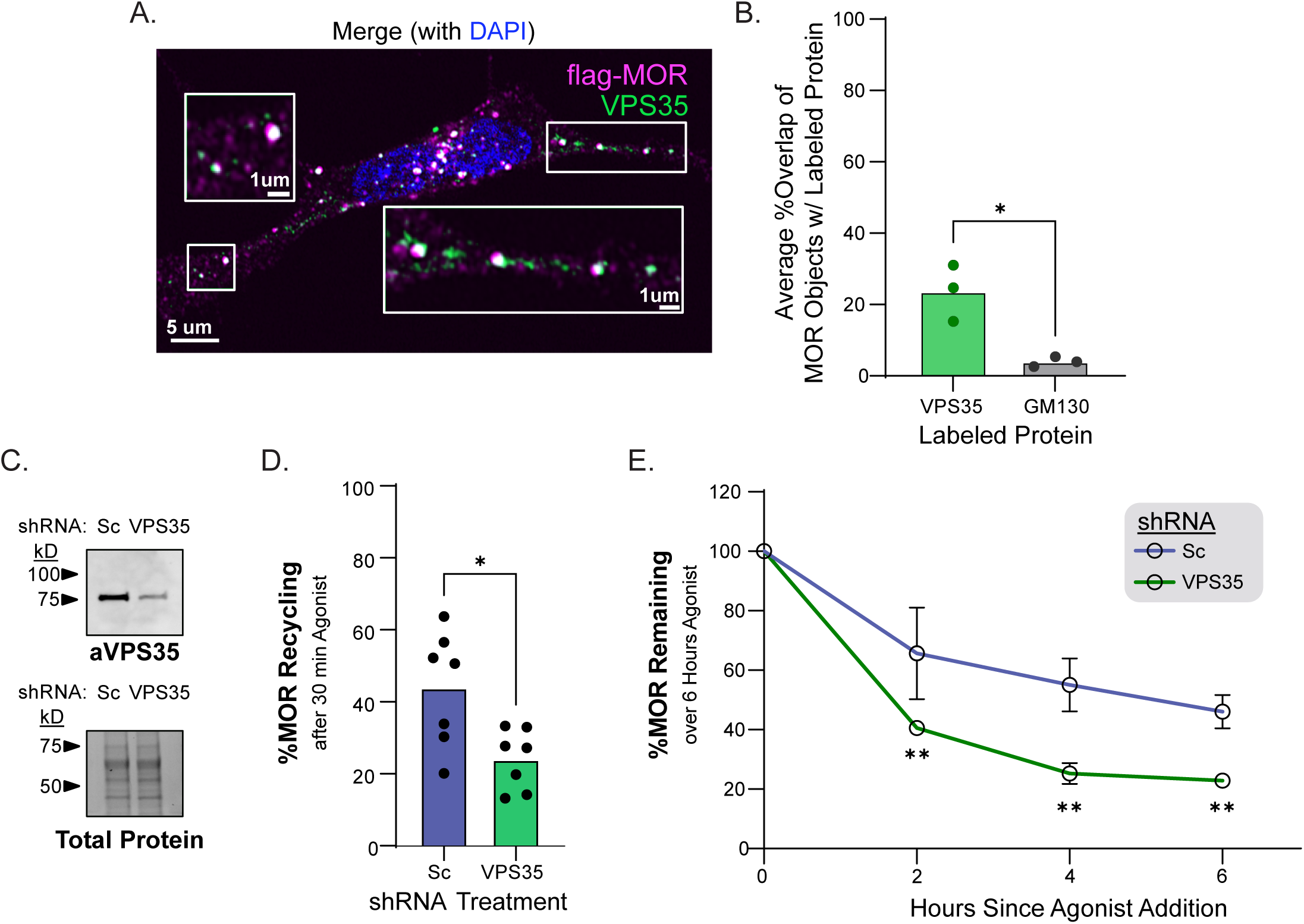
Retromer’s role in MOR recycling is conserved across cell lines. **A.** Confocal images of SH-SY5Y cells stably expressing MOR(WT) and treated with 10μM DAMGO, labeled for anti-FLAG (magenta) and anti-VPS35 (green). Representative image, n=3. **B.** Average percent overlap of all MOR objects with either VPS35 or GM130 (n=3, unpaired two-tailed t-test, p=0.0076). **C.** Western blot for VPS35 and total protein from HEK293 lysates following transduction with either Scramble shRNA or VPS35 shRNA. Representative blot, n=3. **D.** Percent MOR recycling measured by surface labeling of receptors following treatment with 30 minutes 10μM DAMGO to induce internalization and 30 minutes 10μM naloxone to allow for recycling in cells transduced with Sc or VPS35 shRNA. (n=7, paired t-test, p=0.0205 for Sc vs. VPS35). **E.** Percent MOR(WT) remaining following transduction with Sc or VPS35 shRNA and treatment with 0, 2, 4, or 6 hours of 10μM DAMGO followed by the APEX/AUR reaction and normalized to no agonist treatment (n=3, 2 way repeated measures ANOVA with Dunnett’s multiple comparisons, p<0.0001 for time effects, p=0.0368 for shRNA effects, p=0.0054, 0.0022, 0.0081 for Sc vs. VPS35 at 2, 4, and 6 hours respectively). Error bars denote S.D.

To determine the role of Retromer in MOR trafficking, we transduced the MOR(WT) SH-SY5Y line with a previously validated shRNA sequence against VPS35 or a scrambled sequence (Sc). (Choy et al., 2014; Temkin et al., 2017). We found that the shRNA targeting VPS35 resulted in VPS35 knockdown five days after transduction (**Fig. 4C**, **Supp. Fig. 5E**). We then asked how knockdown of VPS35 affected MOR trafficking in SH-SY5Y cells and found loss of Retromer function decreased the ability of MOR to recycle (43.92% vs. 24.05%, p=0.0205 for Sc vs. VPS35) (**Fig. 4D**, **Supp. Fig. 5F**). Finally, we measured the effects of VPS35 knockdown on MOR downregulation over a six-hour period. We found that knockdown of VPS35 significantly increased the rate at which MOR was downregulated (46.01% receptor remaining vs. 22.86% receptor remaining for Sc vs. VPS35 at six hours, p=0.0081) (**Fig. 4E**). Together, these results demonstrate that the role for Retromer function in promoting MOR recycling and opposing opioid-induced MOR downregulation is conserved in human neuronal-derived cells.

### Retromer-dependent trafficking of MOR is contingent on agonist efficacy

Thus far, we focused on the high efficacy peptide agonist DAMGO. DAMGO-stimulated MOR operates similarly to MOR activated by clinically relevant efficacy opioids like fentanyl or methadone (McPherson et al., 2010). Thus, we predicted that fentanyl or methadone-stimulated MOR would also require Retromer function to be protected from opioid-induced downregulation.

Stimulation of HEK293 cells stably expressing MOR(WT) with a saturating dose of fentanyl or methadone induced efficient MOR internalization to a similar level as the peptide agonist DAMGO and, consistent with our hypothesis, recycling of MOR following stimulation with either opioid was Retromer dependent (**Fig. 5A&5B**). We also found that DAMGO, fentanyl, and methadone could all induce downregulation of MOR, and that this downregulation was enhanced upon knockdown of Retromer (**Fig. 5C**). Thus, these data demonstrate that Retromer plays an important role in protecting MOR from opioid-induced downregulation by multiple clinically relevant ligands.

**Figure 5.**
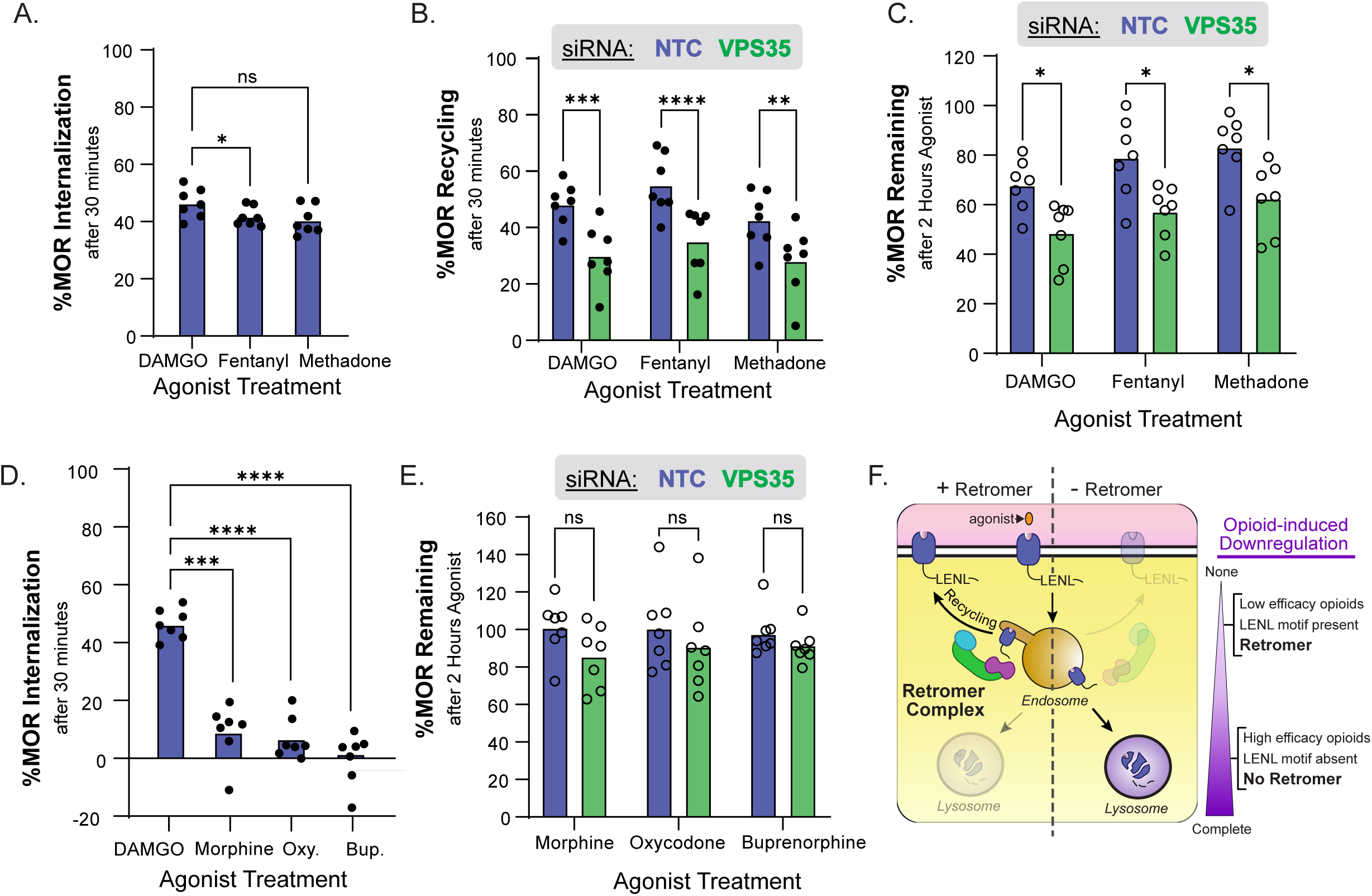
Retromer-dependent trafficking of MOR is contingent on agonist efficacy. **A.** Percent internalization of MOR(WT) in HEK293 cells after 30 minutes of 10μM agonist treatment measured with surface receptor labeling (n=7, repeated measures 1way ANOVA with Dunnett’s multiple comparisons correction, p=0.0375 for agonist effects, p=0.0163 and 0.0709 for DAMGO vs. fentanyl and methadone respectively). **B.** Percent recycling of internalized receptors following 30 minutes of 10uM agonist treatment and 30 minutes of 10μM naloxone treatment measured by surface receptor labeling (n=7, repeated measures 2way ANOVA with Sidak’s multiple comparisons correction, p=0.1866 for agonist effect, p<0.0001 for siRNA effect, p=0.0002, <0.0001, 0.0021 for NTC vs VPS35 for DAMGO, fentanyl, and methadone respectively). **C.** Percent MOR(WT) remaining after two hours of stimulation with 10μM full agonist measured using the GPCR-APEX2/AUR assay following knockdown of VPS35 (n=7, repeated measures 2way ANOVA with Sidak’s multiple comparisons correction, p<0.0001 for agonist effect, p=0.0086 for siRNA effect, p=0.0264, 0.0382 and 0.0411 for NTC vs VPS35 for DAMGO, fentanyl, and methadone respectively). **D.** Same as A but including partial agonists (n=7, repeated measures 1way ANOVA with Dunnett’s multiple comparisons correction, p<0.0001 for agonist effects, p=0.0002, <0.0001, and <0.0001 for DAMGO vs. morphine, oxycodone (Oxy.), and buprenorphine (Bup.) respectively). **E.** Same as C, but using partial agonists (n=7, repeated measures 2way ANOVA with Sidak’s multiple comparisons correction, p<0.8520 for agonist effect, p<0.1998 for siRNA effect, p=0.2655, 0.8305, 0.5681 for NTC vs VPS35 for morphine, oxycodone, and buprenorphine respectively). F. Working model describing how Retromer acts through the LENL recycling motif to protect MOR from agonist-induced downregulation promoted by high efficacy opioids.

Some clinically relevant opioids – such as morphine, oxycodone, and buprenorphine -are lower efficacy MOR agonists. Unlike higher efficacy opioids, lower efficacy opioids cause little, or no, MOR internalization (Arden et al., 1995; Keith et al., 1996), and extensive research suggests that a distinct cellular pathway is involved in regulating MOR when activated by these partial agonists (see discussion) (Johnson et al., 2006; Pena et al., 2018; Melief et al., 2010; Adhikary et al., 2022). Thus, we hypothesized that Retromer’s role in protecting against opioid-induced MOR downregulation would only be observed with high efficacy opioids. Fitting with previous observations, lower efficacy opioids induced much less internalization than the higher efficacy opioids (**Fig. 5D**). Specifically, morphine and oxycodone, but not buprenorphine, induced a small, but measurable, amount of internalization of MOR over 30 minutes (9.12%, 6.883%, and –0.0034% internalization and p=0.0478, 0.0455, and 0.9992 respectively for morphine, oxycodone, and buprenorphine, one-sample t-test against 0). Consistent with this observation, partial agonists induced negligible downregulation of MOR, although we noted a non-significant trend toward enhanced downregulation induced by morphine following VPS35 knockdown (**Fig. 5E**).

Together, these data suggest a working model in which Retromer plays a conserved role in protecting MOR from downregulation in response to high efficacy opioids including fentanyl and methadone—via a previously unrecognized type of Retromer-based recycling motif—by promoting MOR recycling from endosomes (**Fig. 5F**).

## DISCUSSION

Agonist-induced GPCR trafficking plays a critical role in the regulation of many receptors. GPCR recycling from endosomes protects the receptor from rapid agonist-induced downregulation.(Cao et al., 1999; Tanowitz and von Zastrow, 2003; Temkin et al., 2011; Law et al., 2000; Xiong et al., 2016) However, many GPCRs lack consensus recycling motifs, and thus the mechanisms of their trafficking remain unclear (Irannejad and Lobingier, 2022). Here we focused on a GPCR with a non-consensus recycling motif (LENL), MOR (Tanowitz and von Zastrow, 2003). Using a genome-wide screen designed to identify factors that respond specifically to the LENL motif, we identified all three subunits of the endosomal recycling complex Retromer. We showed that the LENL motif is necessary and sufficient to allow Retromer to induce opioid receptor recycling, and that Retromer-dependent recycling protects MOR from downregulation following stimulation by full agonists like fentanyl, methadone, and endogenous-like opioid peptides.

### Implications for recycling pathway diversity in cells

Classically, membrane protein recycling was thought to occur through a sequence-independent “bulk flow” pathway (Mayor et al., 1993). However, it is now clear that many membrane proteins have cis-acting recycling motifs—often in their C-terminal tails—that promote their recycling by binding to endosomal recycling complexes and sorting into endosomal tubules (Puthenveedu et al., 2010; Cullen and Steinberg, 2018). One example is the Retromer complex (VPS35/VPS29/VPS26A), which binds to the consensus sorting motif [W/F/Y]x[L/M/V] (Cullen and Steinberg, 2018; Yong et al., 2022; Harterink et al., 2011; Tabuchi et al., 2010; Seaman, 2007). In the last fifteen years, a number of additional adaptors, cargo binding complexes, and recycling motifs have been found in mammalian cells. These include SNX27/Retromer ([D/E][S/T]xΦ-COOH) (Temkin et al., 2011; Cao et al., 1999; Lauffer et al., 2010; Steinberg et al., 2013; Clairfeuille et al., 2016), SNX17/Retriever ([N][P/X]x[F/Y])(Chen et al., 1990; Böttcher et al., 2012; Butkovič et al., 2024), and SNX-BAR/ESCPE-1 (Φx[F/Y/V]x[F/Y]) (Simonetti et al., 2023, 2017, 2019). However, even this expanded understanding of the diversity of recycling motifs and complexes cannot explain the recycling of many GPCRs and other membrane proteins (Irannejad and Lobingier, 2022).

Here, we demonstrate that the non-consensus mu opioid receptor recycling motif, LENL, is a novel mechanism by which membrane proteins can access the Retromer recycling pathway. It is intriguing to consider whether this novel pathway is unique to MOR, or if LENL-like recycling motifs are also present in other membrane proteins. We used a bioinformatics approach to identify recycling motifs within cytoplasmic C-tails of the human membrane proteome and found that LENL-like LxxL motifs exist in 17% of membrane proteins with cytoplasmic tails, including several already linked to Retromer function including adenylyl cyclase 9 (AC9) and the insulin-responsive glucose transporter GLUT4 (Pan et al., 2017; Yang et al., 2016; Steinberg et al., 2013). Together, our findings here identify a novel type of Retromer recycling motif and raise the potential that a broader portion of the membrane proteome can utilize the Retromer pathway than previously appreciated.

### Implications for Retromer binding mechanisms

An open question is how Retromer recognizes the LENL motif to allow for MOR recycling. The consensus model for Retromer function is that VPS35 acts as a scaffold connecting the membrane proximal cargo-binding subunit VPS26 to the membrane distal regulatory subunit, VPS29 (Kovtun et al., 2018; Baños-Mateos et al., 2019; Leneva et al., 2021; Chandra et al., 2020; Martínez-Núñez and Munson, 2020). A recent structural study has provided further insight into how VPS26 binds cargo containing the consensus motif [W/F/Y]x[L/M/V] (Lucas et al., 2016). This structure resolved an extensive set of contacts between VPS26, the consensus recycling motif from DMT1-II, 551-QPEL**Y**L**L**-557, and Retromer binding protein SNX3. The primary contacts from DMT1-II are L557 dipping into a hydrophobic pocket in VPS26, E553 and Y555 forming a hydrogen bond with SNX3, L554 and L556 forming a bracket around F287 from VPS26, and an extended network of main-chain hydrogen bonds (Lucas et al., 2016). Thus, residues outside the consensus motif (YLL) are important in binding SNX3/Retromer. In this light, it is interesting to note that L554 and L557 in the DMT1-II recycling motif (544-**L**YL**L**-557) —which make key contacts with VPS26—form the LxxL hallmark of the LENL motif in MOR. Furthermore, evidence suggests that the cargo binding pocket in VPS26 can indeed bind peptides that differ from the consensus motif (Lucas et al., 2016; Suzuki et al., 2019). These observations suggest that the current [W/F/Y]x[L/M/V] consensus motif, which was primarily built on analysis of CI-MPR(Seaman, 2007), DMT1-II (Tabuchi et al., 2010), and Wntless (Gasnereau et al., 2011), may only capture one type of sequence that can engage Retromer.

### Other potential MOR trafficking regulators from the genetic screen

The primary finding from the genome-wide screen was that all three subunits of Retromer (VPS35, VPS29, and VPS26A) act through the LENL motif to oppose opioid-induced GPCR downregulation. In alignment with this observation, we previously showed—using APEX2-based proximity proteomics—that Retromer subunits are enriched in the MOR proximal proteome following stimulation with high efficacy agonists (Polacco et al., 2024). In addition to Retromer, we identified several other hits in our genome-wide screen—some specific to DOR(MCT) and others shared with DOR— which likely function in opioid receptor trafficking at the Golgi (RHOBTB3, TRAPPC11), plasma membrane (ARF6, DNM2), and endosomal-lysosomal pathway (GAPVD1, WDR91, RPTOR). Most of these hits represent new proteins potentially involved in MOR trafficking, with the exception of ARF6, which is known to regulate MOR endocytosis and recycling (Claing et al., 2001; Poupart et al., 2007; Donaldson and Jackson, 2011; Macia et al., 2012; Rankovic et al., 2009). We also noticed that several hits only observed in the DOR(MCT-WT) screen were genes which control transcription or translation, which we had initially anticipated would be shared between the two screens. We reason that these hits were identified in one screen, rather than both, because of differences in screen performance (e.g., relative efficacy of the CRISPRi knockdown between the independent cell lines) or design (e.g., seven sub-libraries vs three sub-libraries, 500-coverage vs 300-fold sgRNA coverage).

### Implications for MOR signaling

GPCR signaling and sub-cellular localization are deeply intertwined (Stoeber et al., 2018; Brighton et al., 2024; Vargas et al., 2023). MOR localizes to, and signals from, multiple subcellular compartments including the plasma membrane, endosomes and Golgi (Radoux-Mergault et al., 2023; Stoeber et al., 2018). Our data demonstrate that Retromer function is important in maintaining the distribution of MOR across subcellular compartments in a manner dependent on time and agonist. In this context of shifting subcellular distributions, it is intriguing to consider the multiple—and potentially opposed—ways that Retromer affects MOR signaling. At the plasma membrane, we anticipate that Retromer depletion would cause a decrease in signaling due to inability to reinsert functional receptors following agonist-induced internalization. At the same time, our results suggest that decreased recycling temporarily increases MOR residence time on endosomes, which could potentially lead to increased endosomal signaling. We also anticipate that prolonged agonist exposure would reduce signaling from all compartments by driving MOR downregulation, and that the rate of this process would depend on Retromer function. Thus, a complete analysis of Retromer’s role in MOR signaling will require careful measurements of signaling with high subcellular and temporal resolution.

### Cellular mechanisms regulating MOR function, efficacy, and opioid tolerance

One of the intriguing features of opioids is that the cellular mechanisms which regulate MOR differ when MOR is stimulated by high efficacy opioid agonists compared to lower efficacy opioid agonists. Higher efficacy opioids cause GRK2/3-mediated hierarchical phosphorylation of MOR, efficient beta-arrestin binding, and endocytosis (Williams et al., 2013). Conversely, lower efficacy opioids cause minimal GRK-based phosphorylation of MOR, poor beta-arrestin binding, and little endocytosis (Mann et al., 2015; Miess et al., 2018; Doll et al., 2011, 2012; Lau et al., 2011; Williams et al., 2013). Instead, other kinases including PKC and JNK regulate MOR when stimulated by lower efficacy opioids, presumably in a trafficking independent manner (Bailey et al., 2004; Melief et al., 2010; Johnson et al., 2006; Pena et al., 2018; Bailey et al., 2006). Our work builds on this model by defining the next step in MOR regulation in response to high efficacy opioid agonists: Retromer promotes MOR recycling out of endosomes through the LENL motif and thus protects MOR from opioid-induced downregulation. Additionally, our findings are consistent with the observations that lower efficacy opioids like oxycodone promote minimal MOR endocytosis and suggest Retromer plays a minor role in MOR regulation in these conditions.

A long-standing question in the opioid field is what role these cellular regulatory mechanisms, and in particular MOR trafficking, play in the development of opioid tolerance (Kieffer and Evans, 2002)? This question is particularly interesting because while only high efficacy opioids cause measurable MOR downregulation *in vivo* (Tao et al., 1987; Klee and Streaty, 1974; Stafford et al., 2001; Patel et al., 2002), both low and high efficacy opioids can drive pharmacological tolerance (Hill et al., 2018; Madia et al., 2009; Enquist et al., 2012; Grecksch et al., 2011). Interestingly, several lines of evidence suggest that MOR trafficking contributes to opioid tolerance *in vivo* in contexts where MOR internalization can occur. First, mice expressing mutant MORs—which lack a large part of the receptor C-terminus including the LENL motif—develop tolerance to high efficacy opioids faster than wild-type mice (Enquist et al., 2012). Second, mice expressing mutant MORs which gain the ability to undergo substantial endocytosis in response to morphine develop tolerance more slowly when the LENL motif is present in MOR compared to when it is absent (Enquist et al., 2011, 2012). These studies suggest that MOR endocytosis can promote tolerance but that this is opposed by MOR recycling (Gooding and Whistler, 2024), and future studies will be required to determine if Retromer plays a role in tolerance development.

## ACKNOWLEDGEMENTS

We thank John Williams and members of the Lobingier and Williams labs for their helpful advice and critical feedback on this manuscript. We thank Alexander Dagunts for his assistance with custom software. We thank Mark von Zastrow and Paul Temkin for supplying the VPS35 shRNA constructs. This work was carried out with the help of the Flow Cytometry and Advanced Light Microscopy (SCR_009961) core facility resources. B.T.L was supported by GM137835 and OHSU startup funds. A.D. was supported by T32GM141938. H.A. was supported by T32GM142619.

## METHODS

### Chemicals

DAMGO acetate salt (E7384) and DADLE acetate salt (E7131) were purchased from Sigma-Aldrich. Naloxone hydrochloride (0599) was purchased from Tocris. Fentanyl, methadone, morphine, oxycodone, and buprenorphine were obtained through the National Institute on Drug Abuse Drug Supply Program. All drugs were resuspended at 10mM in double-distilled water and stored as frozen aliquots at -20 degrees C. AMPLEX UltraRed (A36006) was purchased from Thermo Fisher Scientific, resuspended at 10mM in anhydrous DMSO, and stored as frozen aliquots at -20 degrees C. Hydrogen peroxide (30% (wt/wt)) (H1009) was purchased from Sigma-Aldrich, stored at 4 degrees C, and diluted in double-distilled water immediately before use. Sodium ascorbate (A7631) was purchased from Sigma-Aldrich, stored at 4 degrees C, and resuspended in double-distilled water immediately before use. BSA, fatty acid free, (A7030) was purchased from Sigma-Aldrich and resuspended in PBS.

### Antibodies

M1 anti-FLAG (F3040) was purchased from Sigma-Aldrich and used at 1:500 to 1:1000 for immunofluorescence. M1 was conjugated to Alexa Fluor 647 (M1-647) using an amine-reactive labeling kit (A20173) from ThermoFisher Scientific and used at 1:1000-1:2000 for immunofluorescence. Anti-VPS35 (NB100-1397) was purchased from Novus Biologicals and used at 1:500 for immunofluorescence and 1:1000 for western blot. Anti-VPS35L (anti-C16orf62, ab97889) was purchased from Abcam and used at 1:1000 for western blot. Donkey anti-mouse 647 (A31571) and donkey anti-goat 488 (A11055) were purchased from Invitrogen and used at 1:1000 for immunofluorescence. Donkey anti-goat 647 (A32849) was purchased from Invitrogen and used at 1:2500 for western blot. StarBright Blue 700 goat anti-rabbit (12004161) was purchased from Bio-Rad and used at 1:2500 for western blot.

### Complementary DNA constructs

UBC:MORwt-APEX2 is encoded by the construct puDNA5-SSF(signal sequence FLAG)-MORwt-APEX2, which was created by cutting puDNA5 at the NheI and BamHI restriction sites and inserting MORwt, which was amplified by PCR from pSYN-MORwt, and a linker (GGGSGGG) with APEX2, which was encoded by a gBlock. UBC:MOR2ala-APEX2 was created by cutting puDNA5 at the NheI and BamHI restriction sites and inserting MOR2ala and linker-APEX2, which were both amplified by PCR from the UBC:MORwt-APEX2 construct. During PCR amplification, L407 and L410 in the MORwt sequence were replaced with alanines. UBC:DORwt-APEX2 is encoded by puDNA5-SSF-DOR-APEX2 (reference). UBC:DORmct(wt)-APEX2 was created by cutting puDNA5 at the NheI and BamHI restriction sites and inserting the DORwt sequence, which was PCR amplified from pSyn-DORwt-APEX2 (ref) and a sequence containing the last 17 amino acids of MORwt, a linker, and the APEX2 tag, which was encoded by a gBlock. UBC:DORmct(2ala) was created by cutting puDNA5-DORwt with BamHI and inserting the sequence for the last 17 amino acids of MORwt with the two leucines mutated to alanines and a linker, which was encoded by a gBlock, and the sequence for APEX2, which was amplified by PCR from UBC:DORmct(wt). pLenti UBC:MORwt-APEX2 was created by cutting pSYN-MOR-APEX2 at the PacI and XbaI restriction sites to remove the SYN promoter and inserting the UBC promoter, which was PCR amplified from pUBC-MOR-APEX2-Puro. pLenti scramble shRNA and pLenti VPS35 shRNA were gifts from Paul Temkin (Biogen) and Mark von Zastrow (UCSF).

### Cell culture and stable cell line generation

FLP-In-293 (HEK293-FLP, R75007) cells were purchased from Thermo Fisher Scientific and grown in DMEM (11965-092, Thermo Fisher Scientific) supplemented with 10% FBS (Cytiva) at 37 degrees C and 5% CO2. Stable cell lines were created by transiently transfecting either puDNA5-MORwt-APEX2, puDNA5-MOR2ala-APEX2, puDNA5-DORwt-APEX2, puDNA5-DORmct-APEX2, or puDNA5-DORmct2ala-APEX2 alongside pOG44 using Lipofectamine 2000 (11668019, Thermo Fisher Scientific). Transfected cells were selected with 100μg/mL hygromycin (10687010, Thermo Fisher Scientific) and maintained in 50μg/mL hygromycin. For the genome-wide screen, Lenti-X HEK293 cells (632180, Takara Bio) were transiently transfected with pHR-SFFV-dCas9-BFP-KRAB, pVSVG, and psPAX2 with Lipofectamine 2000. The supernatant was collected after 48 hours and filtered through a 0.45-μm PES filter, then incubated overnight with HEK293 cells stably expressing puDNA5-DORmct-APEX2. Cells were double sorted for BFP-dCas9 expression and M1-647 MOR expression. Individual clones were isolated and assessed for dCas9 activity. SH-SY5Y cells were purchased from ATCC and grown in DMEM with 10% FBS at 37 degrees C and 5% CO2. Stable cell lines expressing puDNA5-MORwt-APEX2 were generated using lentiviral transduction as described above. SY5Y cells stably expressing puDNA5-MORwt-APEX2 were transduced with Scramble shRNA-CMV-GFP, or VPS35 shRNA-CMV-GFP packaged into lentivirus.

### Receptor surface expression, internalization, and recycling

HEK293-FLP cells were plated in 12 well plates. 48 hours later, cells were treated with agonist and/or antagonist to determine surface expression, internalization or recycling. The total condition was treated with 10μM naloxone for 30 minutes. The internalization condition was treated with 10μM agonist (DAMGO for MOR cell lines or DADLE for DOR cell lines) for 30 minutes. The recycling condition was treated with 30 minutes of agonist followed by a wash with PBS and 30 minutes of naloxone treatment. Cells were washed once with PBS, lifted with TrypLE Express, and resuspended in PBS with calcium and magnesium supplemented with 1% BSA and 1:1000-1:2000 M1-647. Cells were labeled with M1-647 for 1 hour at 4 degrees C, then washed once and resuspended in PBS with calcium and magnesium and 1% BSA. Cells were analyzed using a CytoFLEX S (Beckman Coulter) using the APC channel (638 nm excitation, 660/20nm emission). Cells were gated for singlets. At least 10,000 singlets were counted for each condition. The geometric mean of the APC channel was used to quantify surface expression of each condition. Internalization was calculated as 1-(internalization geometric mean/total geometric mean), and recycling was calculated as (recycling geometric mean – internalization geometric mean)/(total geometric mean – internalization geometric mean).

SY5Y cells were handled the same way, but after ligand treatment and lifting, were resuspended in PBS with calcium and magnesium supplemented with 1% BSA and 1:1000 M1 and incubated for 1 hour at 4 degrees C. Cells were then washed and resuspended in PBS with calcium and magnesium supplemented with 1% BSA and 1:1000 goat anti-mouse 647. For cells transduced with shRNA, cells were additionally gated for expression of GFP, and 10,000 cells expressing GFP were counted for each condition.

### AMPLEX assay, lysate

Cells were plated in 24 well plates. 48 hours later, cells were stimulated with 10uM agonist (DAMGO or DADLE) for the indicated duration in the legend. Experiments were performed in technical duplicate. Cells were then lysed for three minutes in ice-cold PBS with 0.1% Triton X-100. After lysis, PBS with 0.1% Triton X-100, 100μM AUR, and 200μM H2O2 was added to the cells to provide the necessary substrates for the AMPLEX reaction. 2 minutes later, the reaction was stopped with PBS with 30mM sodium ascorbate. Fluorescence intensity was measured at 555 nm excitation and 610 nm emission on a Spark multimode microplate reader (Tecan). Percent GPCR-APEX2 was calculated as the amount of fluorescence after agonist treatment divided by the amount of fluorescence without agonist treatment multiplied by 100%.

### AMPLEX assay, intact cells

HEK293-FLP cells stably expressing puDNA5-DORmctwt-APEX2 were plated in 24 well plates. 48 hours later, cells were treated with agonist for 2 hours and 15 minutes or left untreated. Cells were then lifted with TrypLE Express and pelleted with DMEM and 10% FBS. Cells were resuspended in ice-cold PBS with 200uM AUR and incubated for five minutes at room temperature, then ten minutes on ice. Next, PBS with 4% (wt/vol) BSA and 100μM H2O2 was added to the cells. Thirty seconds later, the reaction was quenched with 1mM sodium azide. Cells were then washed with PBS with 2% BSA and resuspended in PBS with 1% BSA. Cells were analyzed on a CytoFLEX S using the APC channel.

### Genome-wide CRISPR interference screen

HEK293 cells expressing puDNA5-DORmct-APEX2 and dCas9-BFP-KRAB were transduced with three different CRISPRi sgRNA sublibraries to obtain coverage of the entire human genome. Transduced cells were selected for with 0.75μg/mL puromycin (A1113803, Gibco) 48 hours after transduction. Six days after transduction, the intact cell GPCR-APEX/AUR assay was performed. All samples were pooled and passed through a 40-μm filter, then analyzed on a BD FACSAria II. Cells were gated for singlets, then for BFP positive, and then sorted into the top and bottom quartiles using the APC channel. DNA from these cells was collected with QIAamp DNA Blood Mini kits (51104, Qiagen). sgRNA libraries were prepared using Q5 Hot Start High-Fidelity DNA Polymerase (M0493L, NEB) and barcoding primers. PCR products were purified using QIAquick PCR purification columns (28106, Qiagen) and loaded on 20% TBE gels (EC63155BOX, Thermo Fisher Scientific). A 270 basepair gel was excited from the gel and quantified using a Bioanalyzer (Agilent) and sequenced on an Illumina HiSeq 4000 system (Illumina) using custom primers.

### Bioinformatic analysis of CRISPRi screen and comparison to DORwt-APEX2 screen

Deconvoluted reads from each sublibrary were downloaded from Novogene, aligned to the start of the sgRNAs, and cropped to 38 bp long. Cropped reads for each sublibrary were then loaded into ScreenProcessing (https://github.com/mhorlbeck/ScreenProcessing) python script to count reads of each guide, compute effect sizes by comparing top and bottom quartiles, compile data from multiple individual guides for each gene to compute an average effect and Mann-Whitney P-value of the gene. From negative control guides, pseudogenes were assembled, and hits were determined by thresholding effect and P-value such that less than 10% of hits were pseudogenes.

### Analysis of mouse brain dataset for expression of CRISPRi hits

Single-nuclei RNA sequencing data for each of the 146 hits from the CRISPRi screen, as well as the mu opioid receptor, were downloaded for each cell meta-cluster from www.braincelldata.org. The average percent expression of each hit gene across the top ten MOR-expressing neuron meta-clusters was calculated, and hits were considered expressed in MOR neurons if their average expression was greater than 10%. If a hit was not identified within the dataset, its expression was considered to be 0.

### Small interfering RNA transfections

All siRNAs were purchased from Dharmacon-Horizon Discovery and resuspended in RNase-free water (B-003000-WB) according to the manufacturer’s protocols. The following siRNAs were used: non-targeting control pool (NTC, NC1486135), VPS35 pool (L-010894-00) as well as the four individual siRNAs that make up that pool, VPS29 pool (L-009764-001), VPS26A pool (L-013195-00), Arf6 pool (L-004008-00), VPS35L pool (L-018658-02), and SNX3 pool (L-011521-01). siRNA transfections were performed as “reverse” transfections. 100 pmol of siRNA and 17uL DharmaFECT 1 (T-2001-03, Dharmacon) were incubated for 20 minutes in Opti-MEM (31985070, Thermo Fisher Scientific), then added to a cell suspension and seeded at 40% confluency in a T25 cell culture flask. After 24 hours, cells were split into plates for trafficking experiments. Experiments were conducted 72 hours after transfection.

### Bioinformatics

The entry identifiers for all Swiss-Prot reviewed human proteins tagged with the keyword “Cell Membrane” (KW-1003) were downloaded from the UniProt Knowledgebase. This list of 4019 proteins was uploaded into our custom Motif Searcher application and searched for proteins where the last amino acid in the sequence was within a region with a topological domain annotated (either cytoplasmic or extracellular). This list was downloaded into Microsoft Excel and filtered to exclude proteins with an extracellular annotation resulting in a final list of 2359 proteins with cytoplasmic tails. All remaining searches were performed in this list. The following searches were conducted in the last 100 amino acids of the protein: “L@@L,” “N@@Y,” “[FYM]@[LMV],” and “[GAVCPLIMWF]@@[FYV]@[FY]”, where @ is any amino acid, and any of the bracketed amino acids are accepted within the defined position in the sequence. The PDZ binding motif, “[DE][ST]@[GAVCPLIMFW]” was searched in the last four amino acids of each protein. The resulting lists were downloaded into Microsoft Excel and filtered to exclude motifs found in extracellular regions as well as duplicate proteins from instances where a motif was found multiple times. The final list of unique proteins for each motif was categorized for protein class using the PANTHER database (Thomas et al., 2022).

### Western blot

Cells were lysed in-well with ice-cold RIPA buffer (50mM Tris, 150mM NaCl, 1% Triton X-100, 0.5% sodium deoxycholate, 0.1% SDS, pH 7.4). Halt Protease Inhibitor Cocktail (78430, Thermo Fisher Scientific) was added to the RIPA buffer immediately prior to use. Cell lysates were incubated on ice for ten minutes, sonicated, and centrifuged. The supernatant was added to sample loading buffer with 1% (v/v) 2-mercaptoethanol (1610710, Bio-Rad). Samples were boiled at 95 degrees C for 5 minutes. Proteins were then separated on a Bio-Rad 4-20% Mini-PROTEAN TGX Stain-Free Protein Gel (4568096 or 4568095) in SDS-PAGE running buffer (0.2501 M Tris, 1.924 M glycine, 0.0347 M SDS). Gels were stain-free activated using a Bio-Rad ChemiDoc Imaging System and transferred to nitrocellulose membranes. Membranes were blocked in Bio-Rad Everyblot Blocking Buffer (12010020) for one hour at room temperature, then incubated with primary antibody overnight at 4 degrees C. Blots were washed five times with PBS with 0.1% (v/v) Tween, incubated with secondary antibody for one hour at room temperature, then washed five more times. Blots were imaged on a Bio-Rad ChemiDoc Imaging System. Contrast and brightness were adjusted across the entire uncropped blot using ImageJ-Fiji. Proteins were quantified by normalizing the intensity of the indicated band to the stain-free protein loading control for each lane.

### Fixed imaging

HEK293-FLP cells or SY5Y cells stably expressing puDNA5-MORwt-APEX2 were plated onto poly-L-lysine (P8920, Sigma-Aldrich)-coated coverslips in 24-well plates. 48 hours later, cells were incubated with M1 (1:500) for 30 minutes, washed once with PBS with calcium and magnesium, then treated with 10μM DAMGO for 20 minutes. Cells were then fixed with PBS with 4% (v/v) paraformaldehyde for twenty minutes at room temperature, washed twice with PBS with calcium and magnesium, and blocked and permeabilized with imaging buffer (PBS with calcium and magnesium, 3% (w/v) BSA, and 0.1% (v/v) Triton X-100) for one hour at room temperature. Cells were then incubated with primary antibody in fresh imaging buffer overnight at 4 degrees C. Cells were washed twice with PBS with calcium and magnesium, then incubated with secondary antibodies (1:1000) in imaging buffer for one hour at room temperature. Cells were washed twice with PBS with calcium and magnesium, then mounted on glass slides with ProLong Diamond mounting medium (P36962) and dried overnight. Samples were imaged on a Zeiss LSM 900 with Airyscan 2 with a 63X 1.4NA Plan-Apo lens and Airyscan processed in ZEN Blue software (Zeiss). Single confocal slice images were processed in ImageJ-Fiji.

### Percent overlap analysis of fixed imaging

Airyscan images were converted using Imaris File Converter and 3D surfaces for each channel were produced using the Surfaces tool in Imaris (v10). Surface grain size was set to 0.4 um. Each individual surface object representing an endocytosed receptor (647 channel) was analyzed for its percentage overlap with another surface (488 channel). Three full z-stacks from one slide (technical replicates) were analyzed for three independently prepared slides (biological replicates). Results are displayed as all of surfaces from all of the technical replicates combined, or as the average percent overlap value across the technical replicates.

### Pearson’s correlation coefficient analysis of fixed imaging

Pearson’s correlation coefficient was calculated in Imaris software from three z-stacks per slide (technical replicates) and three independently prepared slides per condition (biological replicates). Images were masked for the 647 channel and automatic thresholding was performed. Each Pearson’s coefficient value displayed represents the average value of three technical replicates.

### Small hairpin RNA transduction

SY5Y cells stably expressing puDNA5-MORwt-APEX2 were transduced with Scramble shRNA-CMV-GFP or VPS35 shRNA-CMV-GFP packaged into lentivirus using LentiX HEK293 cells as described in the cell culture section. Transduced cells were used in experiments five days after transduction to ensure sufficient knockdown and used for up to five passages.

### Statistical analysis and reproducibility

Statistical analysis was performed in Prism (GraphPad) or published software for genomics (ScreenProcessing_v0.1) All experiments except the genome-wide screen include results from at least three biological replicates. Plotted data are represented as individual biological replicates, or as the mean of at least three biological replicates +/-standard deviation, except for supplemental figures 3E and 5C, which show all technical replicates from three biological replicates. The genome-wide screen was performed once across three independent sub-libraries. All measurements were taken from distinct samples, with the exception of the DAMGO internalization data in Figure 5D which is re-plotted from Figure 5A. Statistical testing performed is noted in each figure legend. P values are represented as: ns if P>0.05, * if P<= 0.05, ** if P <= 0.01, *** if P <= 0.001, and **** if P <= 0.0001.

### Software and code

Data were collected with the following software: flow cytometry (Beckman CytExpert, v2.4), plate reader (Tecan Spark Control, v3), western blot (Bio-Rad Image Lab Touch v2.4 and Fiji-ImageJ v1.54f), and microscopy (ZEN v3.5). Data were analyzed with the following software: statistical analysis and graphing (GraphPad Prism v10), flow cytometry (FlowJo v9 or 10), genome-wide screen (open-source custom software ScreenProcessing_v0.1 and RStudio v2023.09.01 build 494), and microscopy (Imaris v10, Fiji-ImageJ v1.54f). The version of the custom code for the Motif Searcher application used in the bioinformatics searches is available on GitHub (https://github.com/dagunts/FASTA-Reader-Public).

---

**Supplemental Figure 1.**
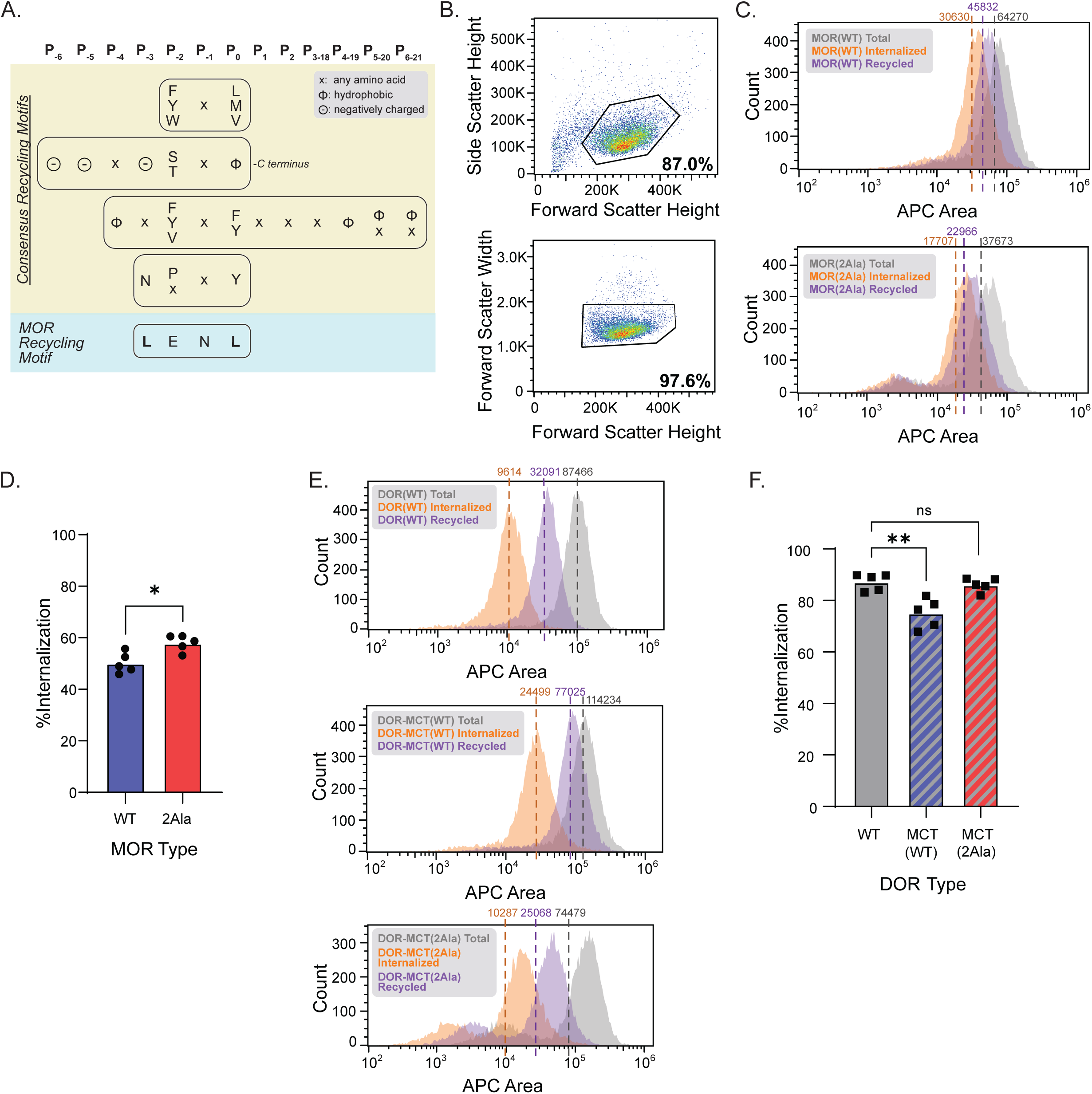
The GPCR-APEX2/AUR downregulation assay captures changes in receptor recycling. **A.** Comparison of known recycling motifs and the MOR recycling motif, adapted from Yong et al 2022. **B.** Example gating scheme for single cells in the flow cytometry recycling assay. **C.** Example histograms of MOR(WT) (top) and MOR(2Ala) (bottom) for cells treated with 10μM naloxone for 30 minutes (total), 10μM DAMGO for 30 minutes (internalized) or 10μM DAMGO for 30 minutes followed by 10μM naloxone for 30 minutes (recycled). Geometric means for each curve are noted. **D.** Percent MOR internalization after 30 minutes of 10μM DAMGO treatment (n=5, two-tailed paired t-test, p=0.0250). **E.** Example histograms of DOR(WT) (top), DOR-MCT(WT) (middle), and DOR-MCT(2Ala) (bottom) for cells treated with 10μM naloxone for 30 minutes (total), 10μM DADLE for 30 minutes (internalized) or 10μM DADLE for 30 minutes followed by 10uM naloxone for 30 minutes (recycled). Geometric means for each curve are noted. **F.** Percent DOR internalization after 30 minutes of 10μM DADLE treatment (n=5, 1way repeated measures ANOVA with Dunnett’s multiple comparisons correction, p=0.0012 for DOR(WT) vs. DOR-MCT(WT), p=0.3087 for DOR(WT) vs. DOR-MCT(2Ala).

**Supplemental Figure 2.**
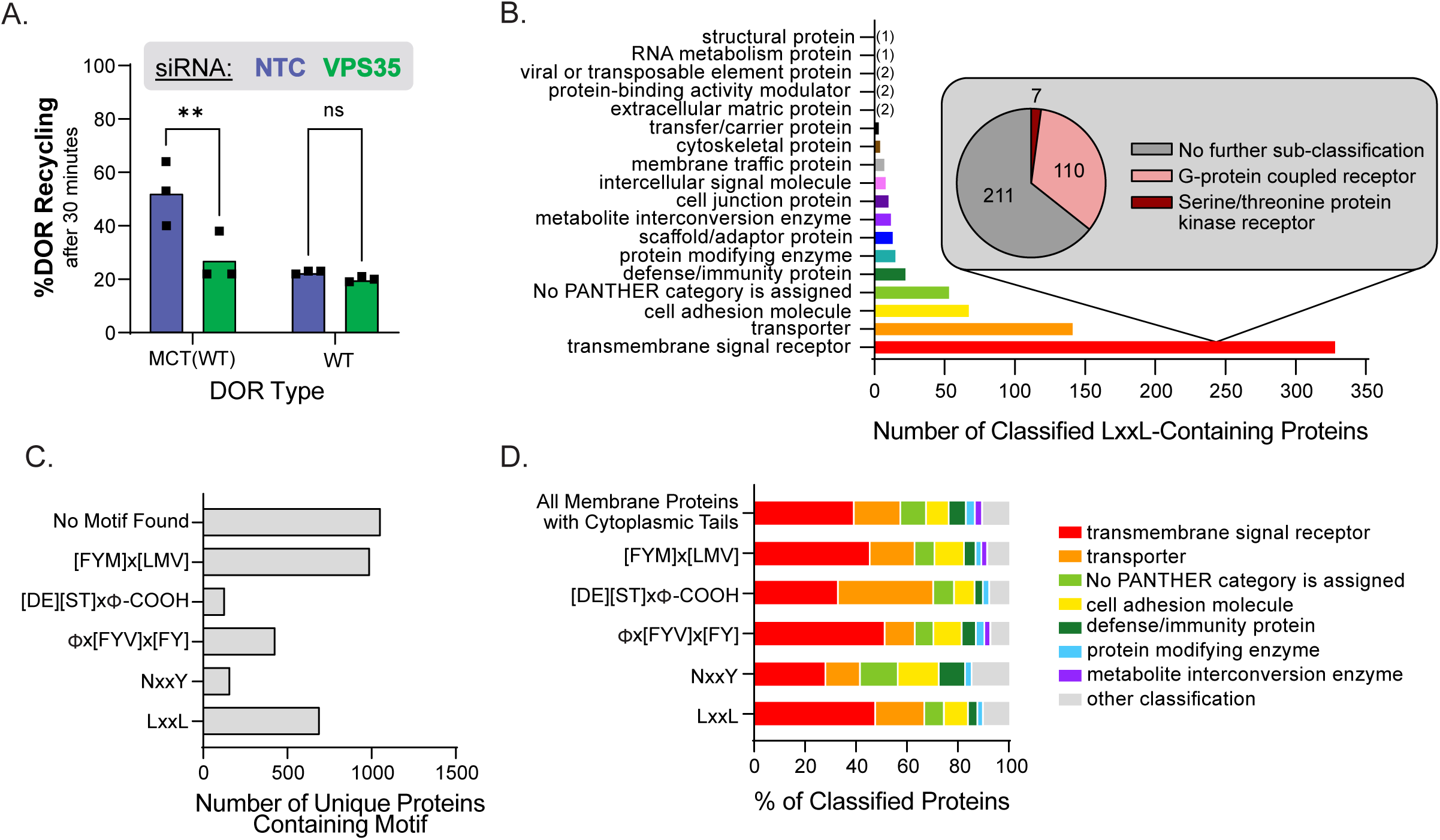
Retromer acts through the LENL recycling motif to oppose lysosomal receptor degradation. **A.** Percent recycling of DOR-MCT(WT) and DOR(WT) in HEK293 cells after VPS35 knock-down and 30 minutes of 10μM DADLE treatment followed by 30 minutes of 10uM naloxone treatment and measured with surface receptor labeling (n=3, repeated measures 2way ANOVA with Sidak’s multiple comparisons correction, p=0.0.351 for DOR type effects for, p=0.0019 for siRNA effects, p=0.0015 for NTC vs. VPS35 for DOR-MCT(WT), p=0.6122 for NTC vs. VPS35 for DOR(WT)). **B.** PANTHER Protein Class analysis of membrane proteins with cytoplasmic C-termini tails containing "LxxL" motifs. **C.** Number of unique membrane proteins with a cytoplasmic tail containing each searched motif. **D.** PANTHER Protein Class analysis of membrane proteins with cytoplasmic tails containing different recycling motifs.

**Supplemental Figure 3.**
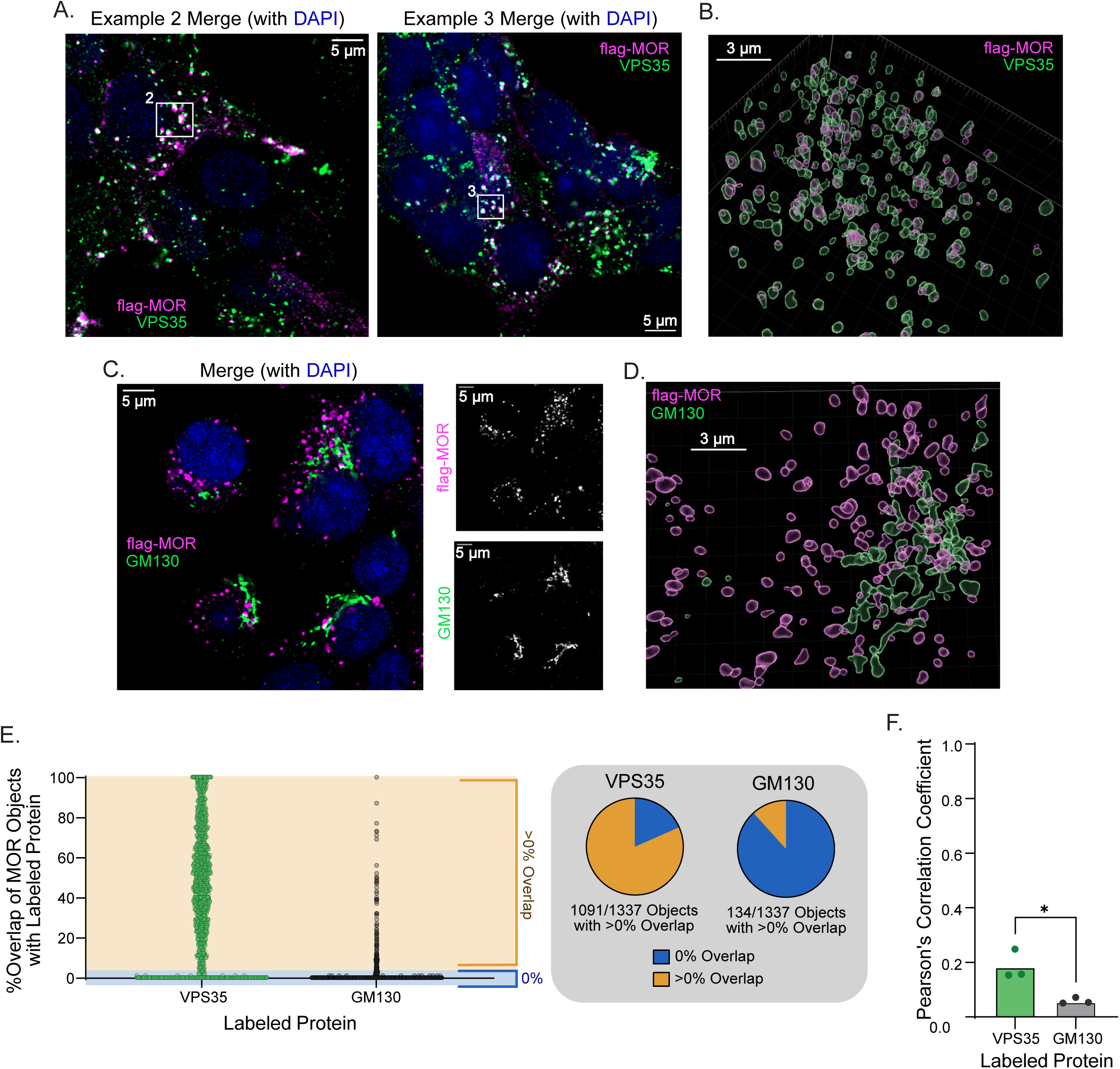
Retromer co-localizes with MOR on endosomes. **A.** Uncropped confocal images of Example 2 and Example 3 (Fig 3A) showing HEK293 cells stably expressing MOR(WT) treated with 10μM DAMGO, labeled for anti-FLAG (magenta) and anti-VPS35 (green). **B.** Example surface rendering of HEK293 cells stably expressing MOR(WT) labeled for anti-FLAG (magenta) and anti-VPS35 (green). **C.** Example confocal image of HEK293 cells stably expressing MOR(WT) labeled for anti-FLAG (magenta) and anti-GM130 (green). **D.** Example surface rendering of HEK293 cells stably expressing MOR(WT) labeled for anti-FLAG (magenta) and anti-GM130 (green) **E.** Percent overlap of individual MOR objects with either VPS35 or GM130. Each point is representative of an individual MOR object from one field from three separate biological replicates. **F.** Pearson’s Correlation Coefficient for co-localization of MOR and VPS35 or GM130 (n=3, unpaired two-tailed t-test, p=0.160).

**Supplemental Figure 4.**
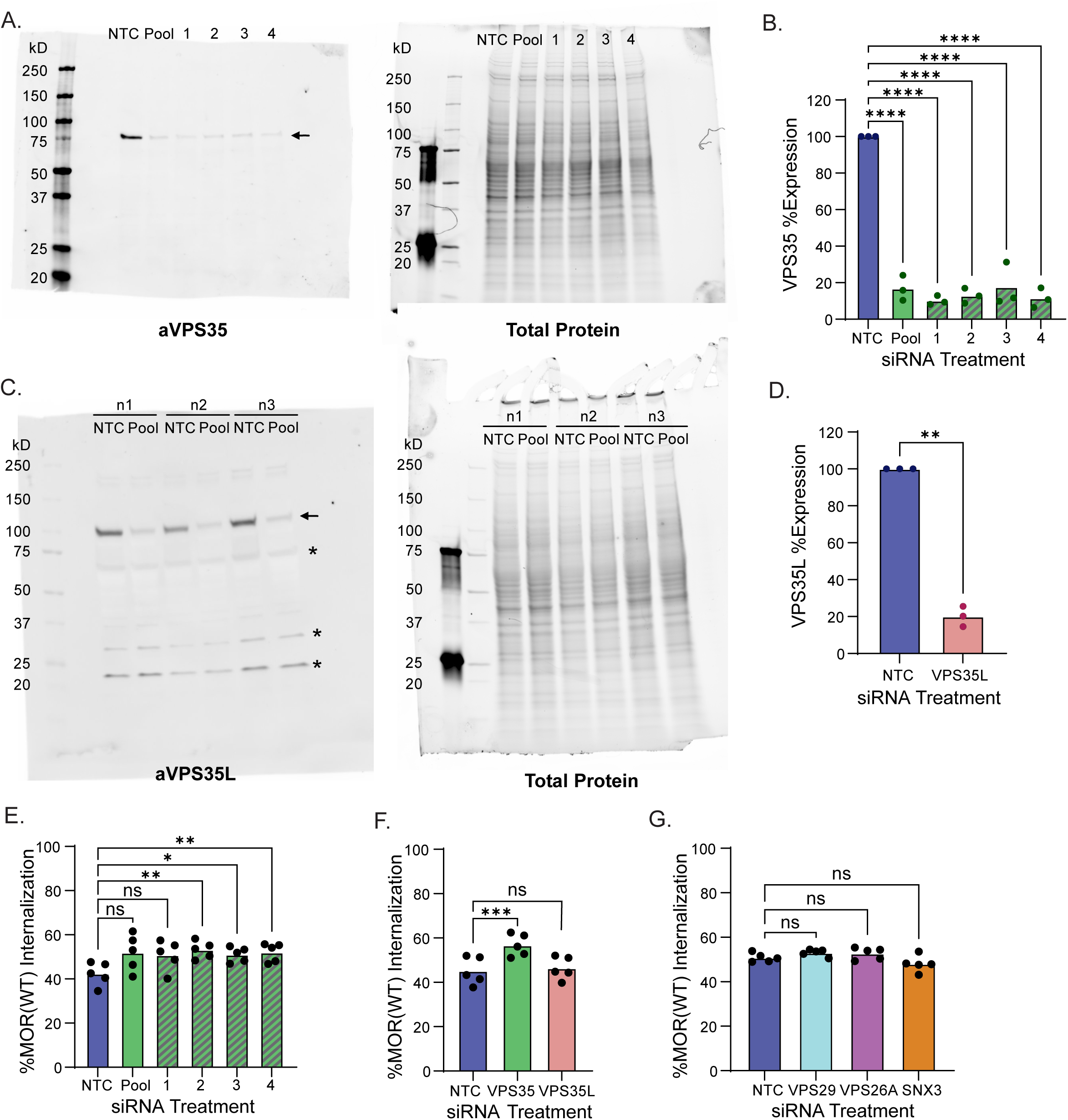
The Retromer complex is required for MOR recycling and resistance to downregulation. **A.** Representative western blot of HEK293 cells stably expressing MOR(WT) treated with either NTC siRNA or siRNA(s) against VPS35. Arrow denotes VPS35. n=3 **B.** Quantification of siRNA knock-down of VPS35 normalized for total protein and to NTC expression (n=3, 1way ANOVA, p<0.0001 for NTC vs. Pool, 1, 2, 3, or 4). **C.** Western blot of HEK293 cells stably expressing MOR(WT) treated with either NTC siRNA or VPS35L siRNA. Arrow denotes VPS35L. Asterisks denote off-target bands. All three biological replicates shown. **D.** Quantification of siRNA knock-down of VPS35L normalized for total protein and to NTC expression (n=3, paired t-test, p=0.0016). **E.** Internalization of MOR(WT) in stably expressing HEK293 cells following siRNA knockdown of VPS35 and treatment with 10μM DAMGO for 30 minutes (n=5, 1way repeated measures ANOVA with Dunnett’s multiple comparisons correction, p=0.0512, 0.1061, 0.0066, 0.0163, and 0.0060 for NTC vs. Pool, 1, 2, 3, and 4 respectively). **F.** Internalization of MOR(WT) in stably expressing HEK293 cells following siRNA knockdown of VPS35 or VPS35L and treatment with 10μM DAMGO for 30 minutes (n=5, 1way repeated measures ANOVA with Dunnett’s multiple comparisons correction, p=0.007 for NTC vs. VPS35 and p= 0.6516 for NTC vs. VPS35L). **G.** Internalization of MOR(WT) in stably expressing HEK293 cells following siRNA knockdown of VPS29, VPS26A, or SNX3 and treatment with 10μM DAMGO for 30 minutes (n=5, 1way repeated measures ANOVA with Dunnett’s multiple comparisons correction, p=0.3343, 0.4733, and 0.3572 for NTC vs. VPS29, VPS26A, and SNX3 respectively).

**Supplemental Figure 5.**
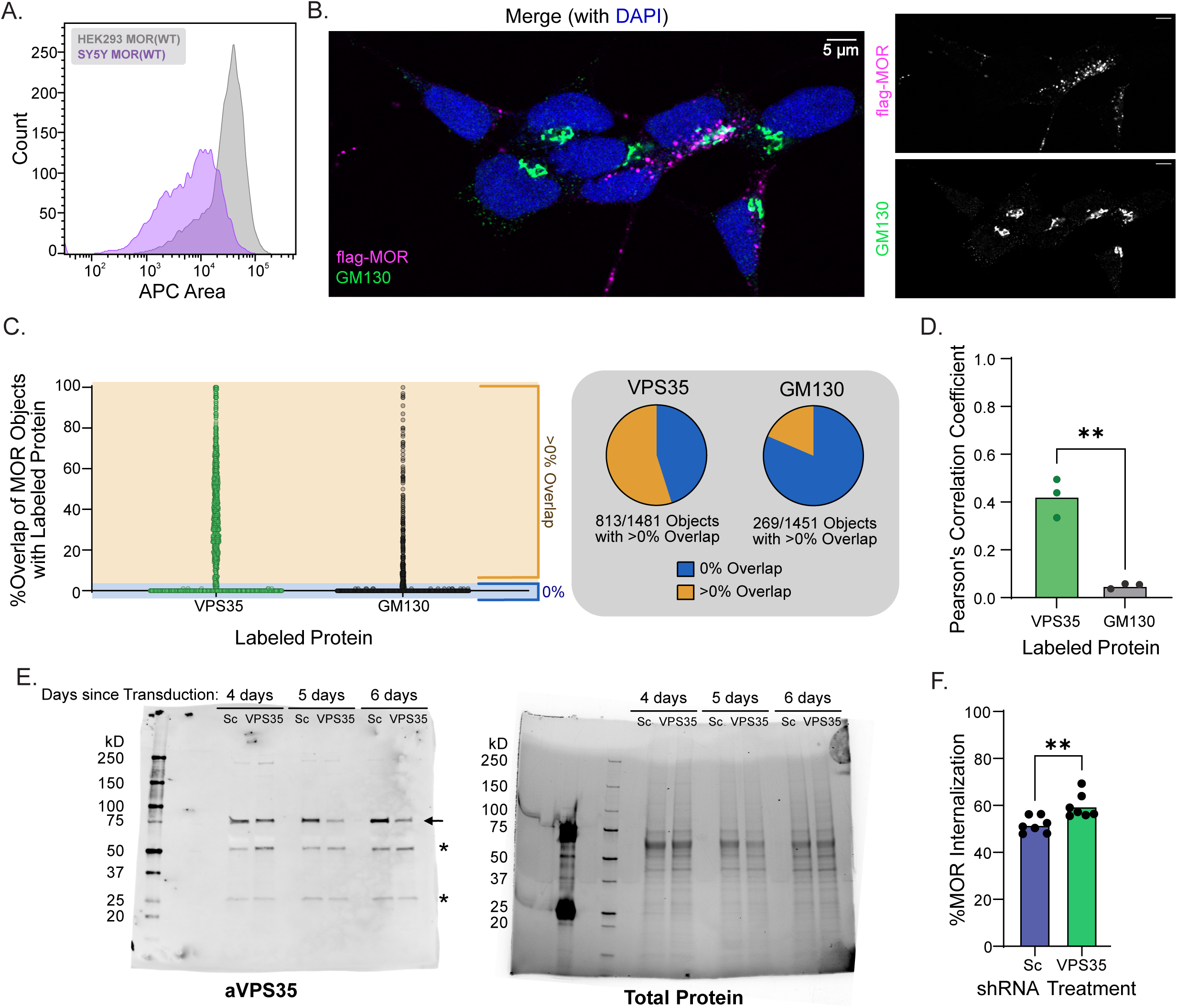
Retromer’s role in MOR recycling is conserved across cell lines. **A.** Flow cytometry histogram comparison of total MOR(WT) expression in the HEK293 and SH-SY5Y cell lines. **B.** Example confocal image of SH-SY5Y cells stably expressing MOR(WT) labeled with anti-flag (magenta) and Golgi labeled with anti-GM130 (green). **C.** Percent overlap of individual MOR objects with either VPS35 or GM130. Each point is representative of an individual MOR object from two fields from three separate biological replicates. **D.** Pearson’s Correlation Coefficient for co-localization of MOR and VPS35 or GM130 (n=3, unpaired two-tailed t-test, p=0.0014). **E.** Representative uncropped western blot for VPS35 and total protein from SH-SY5Y MOR(WT) lysates following transduction with either Scramble shRNA or VPS35 shRNA. Three biological replicates at different timepoints from transduction. Arrow denotes VPS35. Astersisks denote off-target bands **F.** Internalization of MOR(WT) in SY5Y cells transduced with Sc or VPS35 shRNA and treated for 30 minutes with 10μM DAMGO (n=7, paired t-test, p=0.0058 for Sc vs. VPS35).

